# PTPN2 copper-sensing rapidly relays copper level fluctuations into EGFR/CREB activation and associated *CTR1* transcriptional repression

**DOI:** 10.1101/2023.08.29.555401

**Authors:** Matthew O. Ross, Yuan Xie, Ryan C. Owyang, Chang Ye, Olivia N.P. Zbihley, Ruitu Lyu, Tong Wu, Pingluan Wang, Olga Karginova, Olufunmilayo I. Olopade, Minglei Zhao, Chuan He

## Abstract

Fluxes in human intra- and extracellular copper levels recently garnered attention for roles in cellular signaling, including affecting levels of the signaling molecule cyclic adenosine monophosphate (cAMP). We herein applied an unbiased temporal evaluation of the whole-genome transcriptional activities modulated by fluctuations in copper levels to identify the copper sensor proteins responsible for driving these activities. We found that fluctuations in physiologically-relevant copper levels rapidly modulate EGFR/MAPK/ERK signal transduction and activation of the transcription factor cAMP response element-binding protein (CREB). Both intracellular and extracellular assays support Cu^1+^ inhibition of the EGFR-phosphatase PTPN2 (and potentially the homologous PTPN1)–via direct ligation to the PTPN2 active site cysteine side chain–as the underlying mechanism of copper-stimulated EGFR signal transduction activation. Depletion of copper represses this signaling pathway. We additionally show *i*) copper supplementation drives transcriptional repression of the copper importer *CTR1* and *ii*) CREB activity is inversely correlated with *CTR1* expression. In summary, our study reveals PTPN2 as a physiological copper sensor and defines a regulatory mechanism linking feedback control of copper-stimulated MAPK/ERK/CREB-signaling and *CTR1* expression, thereby uncovering a previously unrecognized link between copper levels and cellular signal transduction.

## Introduction

Copper is an essential nutrient in the human diet with prototypical roles as metalloenzyme cofactors; however, excess copper induces cellular (redox) stress via formation of reactive oxygen/nitrogen species. Relatedly, the biochemical mechanisms regulating intracellular copper levels have been extensively characterized. There are two mammalian copper importers; the cuprous importer CTR1 (encoded by the *SLC31A1* gene) – responsible for the vast majority of copper internalization – and the cupric importer DMT1 (encoded by the *SLC11A2* gene). For expediency we hereafter refer to the *SLC31A1* gene and transcript as *CTR1*. Following reduction to Cu^1+^ and import, copper is shuttled throughout the cell by chaperone proteins (e.g. ATOX1) to subcellular locations for metallocofactor assembly or export.

Copper influx and efflux stimulate a wide range of human cellular responses, including cuproplasia (copper-dependent cell growth and proliferation)^1^, cuproptosis^1,2^, neuronal activities^3–5^, cell differentiation^6^, and the epithelial-to-mesenchymal transition in cancer^7^, as well as signaling pathways such as cyclic adenosine monophosphate (cAMP) signaling^8^, MAPK/ERK, JAK-STAT, PI3K/Akt, NF-κB, and the NOD-like receptor protein 3 (NLRP3) inflammasome^9^. Furthermore, copper dyshomeostasis causes human diseases such as Menkes syndrome and Wilson disease, and elevated copper levels induce numerous pro-cancer activities (e.g. angiogenesis, metastasis, and cell proliferation)^10^. Relatedly, serum and tissue copper levels correlate with carcinogenesis and prognosis across diverse cancers^11^. Targeting copper homeostasis pathways has been proposed as promising therapeutic approaches against diverse diseases, including cancer^12,13^.

Across the tree of life, copper stimulates cellular signaling/processes through either indirect mechanisms (change in cellular redox status, generic metal dyshomeostasis response, etc.) or direct binding to proteins. Common bacterial and yeast examples of the latter include copper-sensing transcription factors (TFs), which alter copper-homeostasis protein expression levels to regulate intracellular copper concentrations. In humans, it was recently discovered that copper binding to the enzyme PDE3B inhibits its cAMP-degrading function, leading to elevated levels of the signaling molecule cAMP^8,14^. However, considerably less is known about the transcriptional programs altered by copper levels in humans; whether mammalian intracellular copper levels are even regulated by differential transcriptional/translational expression of copper-homeostasis proteins has been an ongoing debate^15–17^.

In this report, we applied an unbiased temporal evaluation of the whole-genome transcriptional activities stimulated by fluctuations in copper levels under normal cellular growth conditions. We first employed the recently developed kethoxal-assisted single-stranded DNA (ssDNA)-sequencing (KAS-seq) technique^18^, which reports genomic ssDNA regions (corresponding to active transcription), to evaluate the transcriptional programs immediately activated following copper supplementation. By pairing these results with phosphorylation arrays to define copper-stimulated signal transduction pathways, as well as transcriptomic, molecular biological, and biochemical investigations, we produced strong evidence that low level copper supplementation causes Cu^1+^ binding to–and inactivation of–the protein tyrosine phosphatase (PTP) PTPN2 at the active site cysteine to drive activation of EGFR signal transduction, leading to MAPK/ERK/CREB activation. Moreover, we show copper supplementation drives *CTR1* repression, and that CREB activity is inversely correlated with *CTR1* expression. We propose this pathway likely serves as a feedback response to balance copper-uptake and attenuate copper-stimulated cAMP signaling, working in concert with other established mechanisms like copper-stimulated CTR1 protein degradation^19^. Thus, copper-/CREB-stimulated *CTR1* transcriptional repression may drive the previously reported ∼70% decrease in *CTR1* transcript levels following sustained activation of CREB and intracellular copper accumulation during neuronal differentiation^6,20–23^.

## Results

KAS-seq provides a genomic method that enables temporal profiling of the whole-genome transcriptional activity to dissect early responsive genes upon cellular stimulation or signaling. To study copper-induced signaling we applied KAS-seq to evaluate the genomic responses of A549 cells (non-small cell lung adenocarcinoma) to 10 µM CuCl_2_ (low Cu) or 20-30 µM CuCl_2_ (high Cu) supplemented growth media for 15 minutes, 2 hours, and 4 hours, relative to unsupplemented cells (Fig. 1A). For reference, healthy serum copper levels are ∼10-30 µM (of which ∼25% is readily exchangeable^24^), while up to ∼3-fold increases have been observed in cancer patients^11^. Through KAS-seq we captured the sequence of copper-stimulated transcriptional responses (Fig. 1B, SI Appendix Figs. S1-2), including the immediately-activated genes first induced by copper stimulation. Examining the gene body differential ssDNA levels revealed three genes significantly activated by both low and high copper incubation conditions: *EGR1* and *NR4A1* (15 min post-copper) as well as *CXCL2* (2 hours post-copper), all of which are established CREB1 target genes^25^ and are induced by the MAPK/ERK signal transduction pathway^26^.

**Figure 1.**
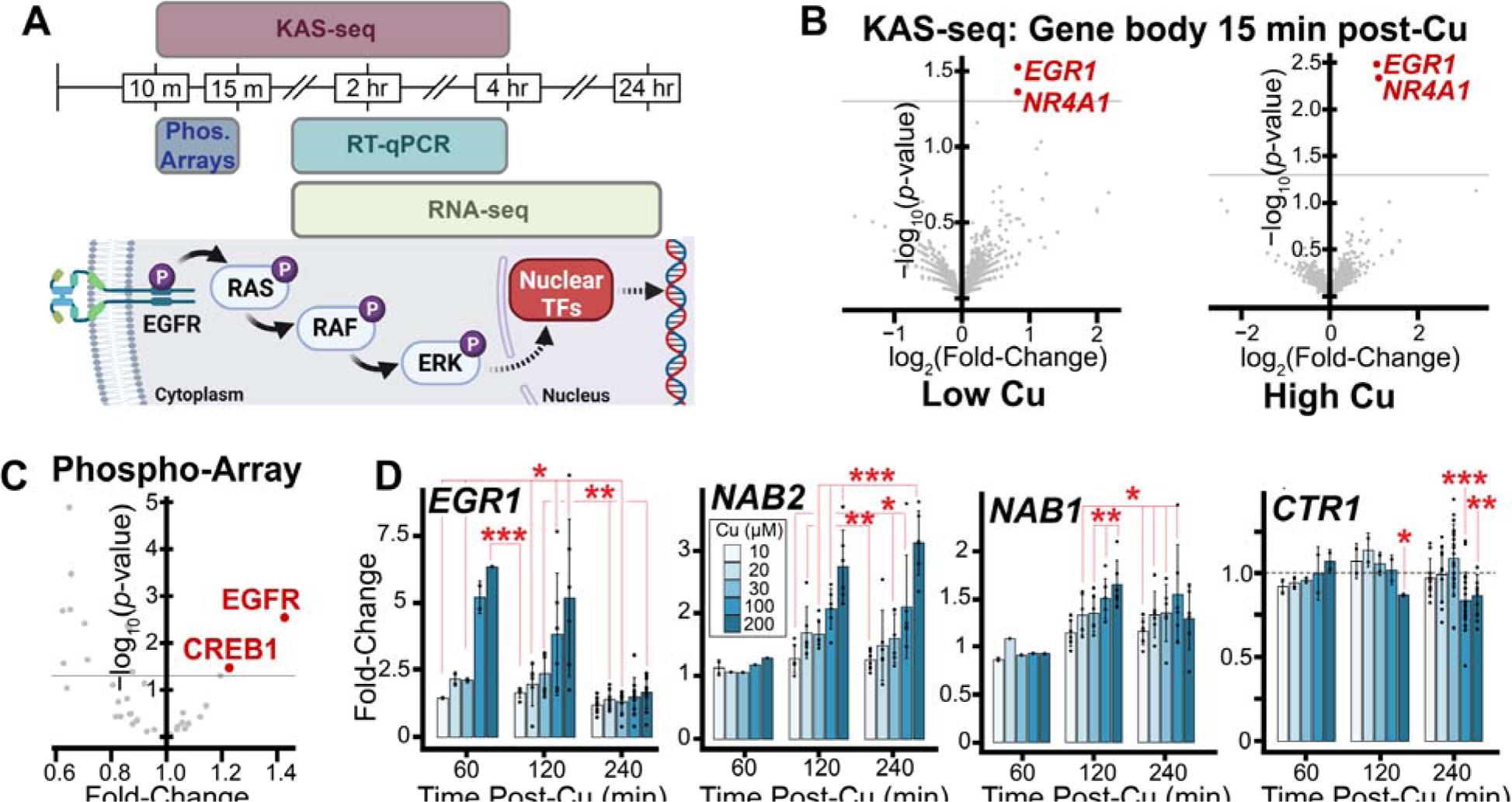
EGFR/MAPK/ERK signaling and associated gene regulation. (*A, Top*) Schematic of the various techniques applied in this study to evaluate copper-stimulated transcriptional/genomic dynamics, with (*Bottom*) the canonical EGFR/MAPK/ERK signaling pathway. Figure created with BioRender.com. (*B*) A549 KAS-seq gene body volcano plots (Cu-supplemented vs. unsupplemented) 15 min post-cell treatment. Significantly upregulated genes in red; horizontal gray lines denote *p*-value = 0.05; *n* = 2 biological replicates for each condition. Low Cu and High Cu conditions correspond to 10 and 20 µM CuCl_2_ supplementation, respectively. (*C*) Changes in HEK 293T phosphorylation and/or total protein levels from proteome profiler phospho-kinase antibody arrays following 10 min, 20 µM CuCl_2_ treatment (vs. untreated). *n* = 3 biological replicates for each condition. (*D*) RNA-qPCR fold-changes of select transcripts following copper supplementation for indicated times into A549 cells. Data points correspond to biological replicates, error bars denote standard deviation, *p*-values calculated as paired two-tailed *t*-test. Sample sizes were as follows for each transcript at 60 min, 120 min, and 240 min post-Cu; *EGR1* (*n* = 2, 6, 10), *NAB1* (*n* = 2, 6, 6), *NAB2* (*n* = 2, 6, 6), *CTR1* (*n* = 2, 2, 10-22).

The observed MAPK/ERK pathway activation by copper led us to evaluate changes in phosphorylation and total protein levels among common signal transduction proteins via proteome profiler phospho-kinase antibody arrays. Because A549 cells express mutant hyperactive KRAS and overexpress EGFR, which could obfuscate detection of copper-stimulated MAPK/ERK activation, we selected a non-cancerous cell line for further validation and investigation. HEK 293T cells were briefly (3 hour) serum-starved to attenuate serum-responsive signaling, then were supplemented with 20 µM CuCl_2_. 10 minutes post-copper supplementation, cells were lysed and evaluated via the array relative to unsupplemented cell lysates (SI Appendix Fig. S3). AKT phosphorylation was strongly, significantly diminished in response to copper treatment – interestingly, the same decrease in AKT signaling was observed when glial cells were treated with a cell-permeable small molecule copper chelate^27^. There were two significantly activated protein phosphorylation sites: EGFR Y-1086 and CREB1 S-133 (Fig. 1C).

Phosphorylation of EGFR Y-1086 (as well as Y-1068) activates MAPK/ERK signaling^28^, which subsequently drives phosphorylation of the nuclear TF CREB1 S-133 site^29^ (among other TFs). The KAS-seq and phosphorylation array results collectively indicate that low level copper supplementation drives EGFR activation to initiate copper-stimulated MAPK/ERK signaling, followed by downstream CREB1 activation and transcriptional responses.

To corroborate the transcriptional activities reported by KAS-seq and provide more quantitative assessment of this potential copper-induced MAPK/ERK signal transduction, we performed RNA-qPCR to analyze copper homeostasis (*ATOX1* and *CTR1*) and MAPK/ERK-induced (*EGR1*, *NAB1*, and *NAB2*) transcript expression levels from cells supplemented with 0-200 µM CuCl_2_. As expected, *ATOX1* expression levels were unresponsive to copper supplementation (SI Appendix Fig. S4). We measured a dose-dependent increase in *EGR1* expression 1 hour post-copper supplementation, followed by increased *NAB1* and *NAB2* expression 1-3 hours later (Fig. 1D). *NAB1*/*2* expression is known to be strongly induced 1 hour after MAPK/ERK-induced *EGR1* expression^30,31^. *CTR1* was repressed 4 hours after ≥100 µM copper supplementation, consistent with prior RNase protection experiments^17^. We subsequently performed RNA-seq and confirmed that these copper-stimulated changes in *NAB1*, *NAB2*, and *CTR1* expression levels were again detected 4 hours post-copper supplementation (SI Appendix Fig. S5). Pre-treatment of A549 cells with EGFR-inhibitors (2 µM gefitinib or 80 µg/mL cetuximab) followed by copper supplementation and RNA-qPCR validated copper-stimulation of EGFR signaling as the driver of increased *EGR1* expression and *CTR1* repression (SI Appendix Fig. S6).

Our observation of copper-stimulated EGFR activation suggested that MAPK/ERK-downstream TFs may be responsible for *CTR1* transcriptional repression. As our results defined copper supplementation drives CREB phosphorylation – and given the aforementioned role of copper in cAMP signaling – we evaluated CREB as the candidate copper-stimulated *CTR1*-repressor TF. We analyzed previous chromatin immunoprecipitation-sequencing (ChIP-seq) experiments, and found significant CREB1 binding to the *CTR1* promoter in the majority of human cell lines in the ChIP-Atlas as well as in the CREB Target Gene Database^32–34^ (Fig. 2A, SI Appendix Fig. S7, Table S1). The *CTR1* promoter *CRE* (cAMP response element) does not contain a proximal TATA box (SI Appendix Table S1), the presence of which is generally associated with CREB-induced gene expression^35^, consistent with a role of CREB-induced *CTR1* repression.

**Figure 2.**
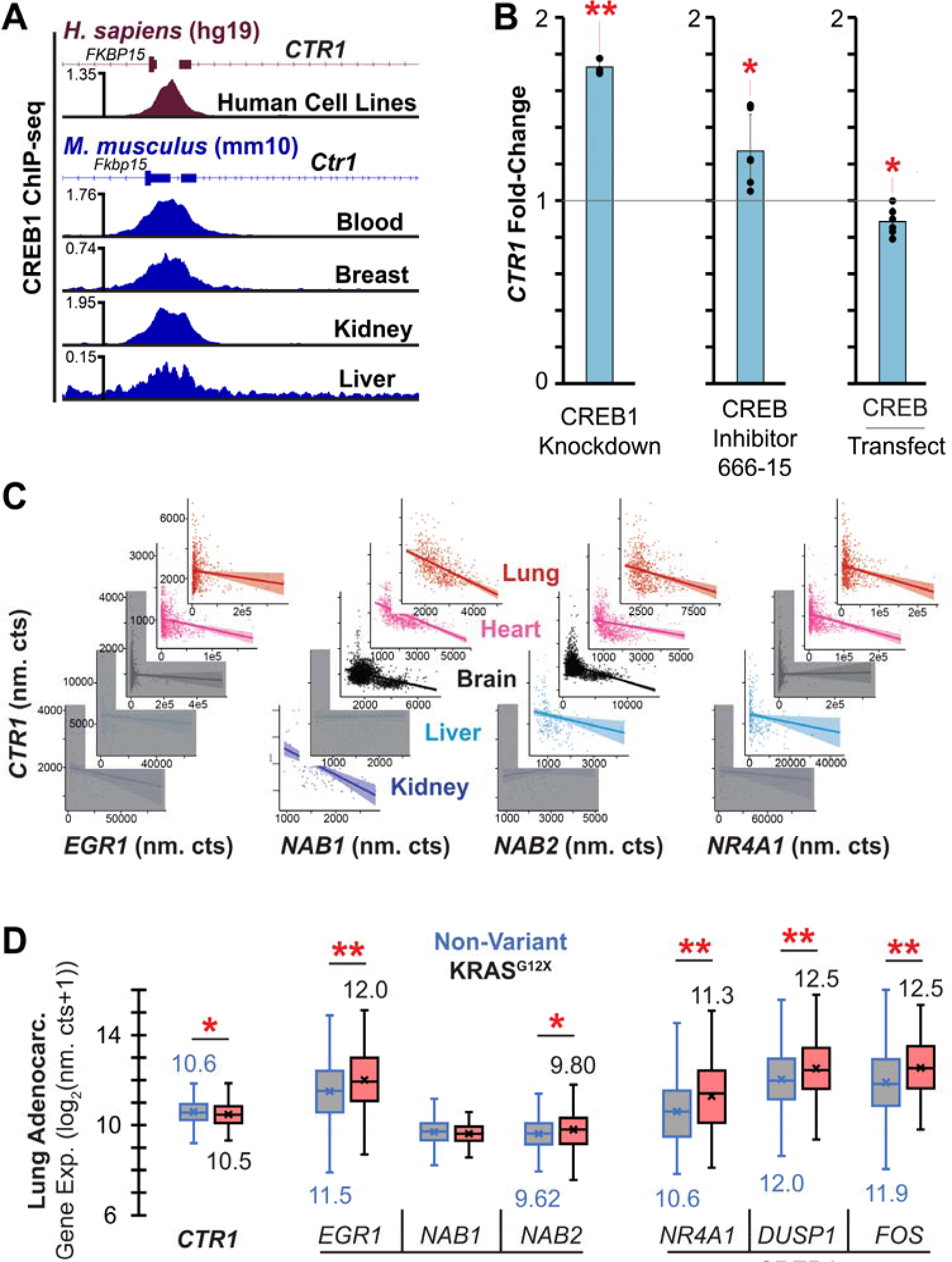
Transcriptomic correlations of MAPK/ERK/CREB1 activation and *CTR1* repression. (*A*) Mean enrichment of CREB1 ChIP-seq experiments generated from ChIP-Atlas *Homo sapiens* cell lines (*Top*) and *Mus musculus* tissues and blood (*Bottom*). Individual ChIP-seq experiments presented in SI Appendix Fig. S7. (*B*) A549 *CTR1* transcript levels following modulation of CREB activity levels. (*Left*) 24 hours post-CREB1-knockdown, relative to cells transfected with negative control (non-targeting) siRNA. *n* = 3 biological replicates for each condition. (*Middle*) 24 hours after 2.56 µM CREB inhibitor 666-15 treatment relative to cells treated with DMSO. *n* = 6 biological replicates for each condition. (*Right*) 24 hours after transfection of a pCMV-CREB1 overexpression vector, relative to transfection with a dsRed-EGFP plasmid. *n* = 6 biological replicates. (*C*) Correlations of MAPK/ERK/CREB1-stimulated transcript expression levels with *CTR1* levels from healthy human tissue (GTEx) transcriptomic data. Correlation line shadow corresponds to 95% confidence interval. Not significant correlations (*p*-value > 0.05) covered in gray boxes. Fit statistics provided in SI Appendix Table S3. *n* = 578, 861, 2642, 226, 89 GTEx RNA-seq samples for lung, heart, brain, liver, and kidney tissues, respectively. (*D*) Boxplots generated from TCGA gene expression data of select MAPK/ERK/CREB1-stimulated transcripts, comparing MAPK/ERK activating KRAS^G12X^ mutant relative to non-variant lung adenocarcinoma transcript levels. Box plot values defined as follows: box boundaries (median of first quartile to median of third quartile) and central horizontal line (median of all expression values), numbers below/above individual box plots denote mean expression value of that dataset (shown on the plot as an “x”), whisker boundaries define 1.5 times the interquartile range; outlier data points omitted for visual clarity. n = 362 and 131 samples for KRAS non-variant and KRAS^G12X^ samples, respectively.

We then sought direct experimental evidence of CREB-induced *CTR1* repression. We first evaluated *CTR1* levels in A549 cells via RNA-qPCR following *CREB1*-siRNA transfection (Fig. 2B, knockdown validation provided in SI Appendix Fig. S8); *CREB1* knockdown significantly increased *CTR1* expression. Notably, the same results were also observed in a *CREB1*-knockdown RNA-seq study in K562 (leukemia) cells (SI Appendix Table S2)^36^. We next assessed chemical inhibition of CREB1 via supplementation of the small molecule CREB inhibitor 666-15, and similarly found *CTR1* levels were significantly increased (Fig. 2B). By contrast, we found cells transfected with a pCMV vector driving constitutive *CREB* expression significantly repressed *CTR1* levels (Fig. 2B). Collectively, these results revealed CREB as the downstream copper-stimulated repressor TF of *CTR1*.

To demonstrate physiological relevance of this pathway, we next sought *in vivo* evidence of *CTR1* repression via CREB. We plotted mean ChIP-signal enrichments from mouse tissue and blood CREB1 ChIP-seq experiments in the ChIP-Atlas (Fig. 2A, SI Appendix Fig. S7). These ChIP-seq results confirmed binding of CREB1 at the *Ctr1* promoter inside living mammals. Relatedly, a prior study found mice injected with FGF23 (a MAPK/ERK activator kidney hormone) exhibited pronounced renal repression of *Ctr1*^37^.

We next evaluated correlations between *CTR1* levels with expression levels of MAPK/ERK responsive genes (*EGR1*, *NAB1*, and *NAB2*) and the CREB1 target gene *NR4A1* across the five vital organs in healthy human tissues using Genotype-Tissue Expression (GTEx) RNA-seq datasets. We assessed whether activation of the MAPK/ERK/CREB1 signaling, as inferred by increased *EGR1*, *NAB1*, *NAB2*, and *NR4A1* expression levels, is associated with repression of *CTR1*. In all tissues, transcript correlations with *CTR1* were negative or statistically insignificant (Fig. 2C, SI Appendix Table S3). As *EGR1*, *NAB2*, and *NR4A1* expression tend to be transient and stimulus-induced^38,39^, the observed inter-tissue-conserved negative correlations support stimulus-responsive *CTR1* repression by CREB1 *in vivo*. Note that *CREB1* levels are also negatively correlated with *CTR1* levels across the aforementioned tissues (excluding the non-significant correlation in the heart tissue, SI Appendix Fig. S9).

For further *in vivo* evidence, and given the important roles of EGFR/MAPK/ERK/CREB1 in lung adenocarcinoma, we evaluated the Cancer Genome Atlas (TCGA) lung adenocarcinoma RNA-seq results, comparing MAPK/ERK-activating KRAS^G12X^ mutant (X is A, C, D, S, or V) with non-variant samples. As expected, MAPK/ERK-stimulated and CREB1-target gene transcripts were significantly increased while *CTR1* levels were significantly decreased in KRAS^G12X^ mutant relative to non-variant samples (Fig. 2D). Collectively, our results provide strong evidence that *CTR1* expression levels are repressed by CREB.

To investigate this newly discovered copper-induced regulatory pathway, as well as demonstrate reversibility of the copper-stimulated CREB1 TF-activation response, we next evaluated differences in gene expression across the transcriptome induced by copper-depletion (via supplementation of tetrathiomolybdate (TM)) or copper supplementation using RNA-seq. Note: TM was chosen as a copper-chelator (as opposed to Cu^1+^/Cu^2+^-specific, or intra-/extracellular specific) as a way to sequester both extra- and intracellular copper. We found a close correspondence between transcriptomic responses 2 hours post-copper supplementation and 24 hours post-TM supplementation (Fig. 3A-B). Pathway and process enrichment analysis of differentially expressed transcripts (FDR-q < 0.05) post-treatment corroborated known copper-stimulated pathways, supporting that these responses are regulated at the transcriptional level. Four ontologies of particular relevance were enriched in both the TM- and copper-supplemented differentially expressed genes (that is, transcripts significantly up-or downregulated, Fig. 3C-D, SI Appendix Table S4-S5): i) regulation of (MAP) kinase activity, ii) response to growth factor (including TGF-β – an EGFR transactivator^40^ – response ontologies), as well as the related iii) epithelial cell differentiation and iv) cell/tube morphogenesis. Moreover, consistent with the known role of copper in stimulating cellular differentiation, evaluation of transcripts significantly repressed in response to TM also revealed enrichment of terms iii) and iv) (SI Appendix Fig. S10). Of further particular note, the copper-treated dataset revealed enrichment of the response to cAMP ontology, consistent with a role of copper in activating CREB.

**Figure 3.**
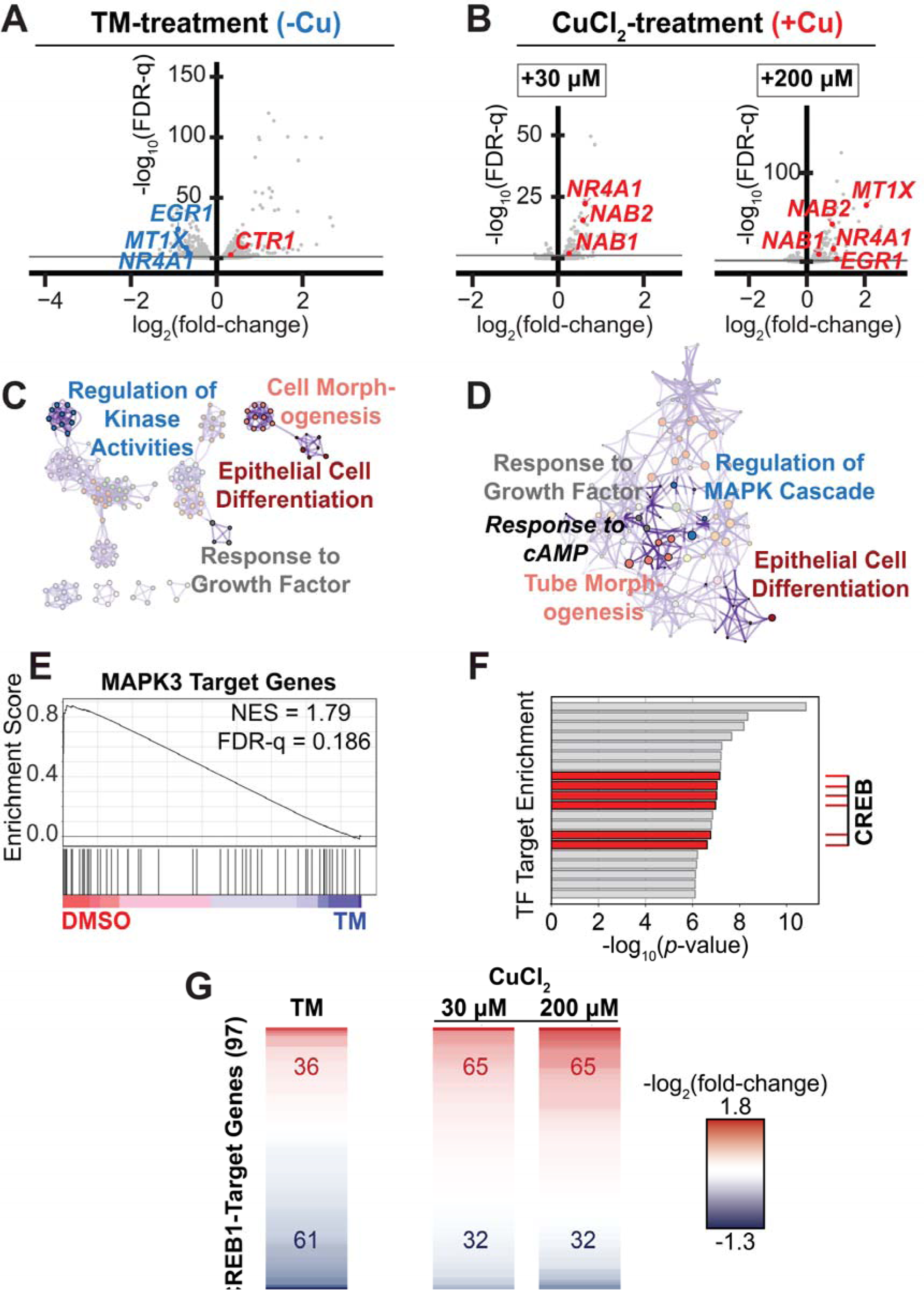
Transcriptomic changes induced by altered copper levels and/or MAPK/ERK/CREB1 signaling. RNA-seq volcano plots depicting differential expression for (*A*) MDA-MB-468 cells treated with TM for 24 hours (relative to DMSO-treated) and (*B*) A549 cells treated with CuCl_2_ (30 µM or 200 µM) for 2 hours (relative to unsupplemented). For visual clarity, outlier datapoints omitted from volcano plots; full plots shown in SI Appendix Fig. S11. As expected, *MT1X* expression levels (which are correlated to intracellular copper levels following copper supplementation^65,66^) were decreased following TM treatment and increased following copper treatment. FDR-q is the false discovery rate-adjusted *p*-value. Gray lines denote FDR-q = 0.05. *n* = 2 (MDA-MB-468) or 3 (A549) biological replicates for each condition. Pathway and process enrichment for significantly differentially expressed transcripts following (C) TM or (D) 30 µM CuCl_2_ treatment. All enriched ontologies provided in SI Appendix Tables S4-5. (E) Transcription factor target gene set enrichment analysis (GSEA) depicts the activation of MAPK3 (a.k.a. ERK1) target genes in DMSO-treated relative to TM-treated cells; NES is normalized enrichment score. (F) TF target enrichment generated from significantly differentially expressed transcripts post-2 hour, 30 µM CuCl_2_ treatment. Multiple CREB hits correspond to different CREB target gene lists. Fully labeled TF target enrichment presented in SI Appendix Fig. S19. (G) Heatmaps depicting differential expression of CREB1-target genes (97 assessed in total) following TM- or CuCl_2_-treatment. Numbers in red and blue denote upregulated and downregulated CREB1-target genes, respectively. Gene list with corresponding expression values presented in SI Appendix Fig. S20. Note: the figure is intended to provide a general assessment of the treatment effects on CREB1-target gene expression – discrete lines (corresponding to individual transcripts) should not be compared across conditions and do not necessarily correspond to the same transcript.

While MAPK3 (a.k.a. ERK1) target genes were significantly depleted upon TM treatment, CREB target genes were significantly enriched in the copper-supplemented dataset (Fig. 3E-F). Indeed, copper generally stimulated activation of known CREB1-target genes, whereas TM generally repressed them (Fig. 3G). Finally, *CTR1* levels were significantly increased following TM treatment, further confirming that decreasing bioavailable extracellular copper drives *CTR1* expression (Fig. 3A and SI Appendix Fig. S11).

To probe the molecular and biochemical mechanism underlying copper-stimulated EGFR activation, we next applied anti-EGFR phosphorylation arrays to briefly serum-starved (∼3 hours) A549 cell lysates generated from cells supplemented with 20 µM CuCl_2_ for 15 minutes, relative to unsupplemented cell lysates. An unambiguous, reproducible ∼50% decrease in pan-EGFR (total EGFR) levels in response to copper was measured (SI Appendix Fig. S12). Rapid (sub-hour-timescale) degradation of EGFR is an indicator of a specific interaction with EGFR, generally a conformational change; in contrast to the slow degradation induced by stress conditions^41^. Note that *EGR1* expression was activated by copper supplementation in cells pre-treated with metalloproteinase inhibitor to a similar magnitude as non-pre-treated cells (SI Appendix Fig. S13), excluding the possibility copper supplementation leads to activation of metalloproteinases which then cleave pro-growth factors and activate EGFR.

Given the rapid EGFR degradation upon copper supplementation, it was important to evaluate the possibility that extracellular Cu^2+^ may drive directly bind EGFR to activate signaling. To probe Cu^2+^-binding by EGFR, we expressed and purified soluble EGFR-extracellular domain (sEGFR; residues 1-642). While we confirmed sEGFR bound Cu^2+^ with a physiologically-relevant K_D_ (200-700 nM; SI Appendix Fig. S14), and formed a well-structured Cu^2+^ binding site as assessed by EPR (SI Appendix Fig. S15); no Cu^2+^-driven dimerization (as required for ligand-dependent EGFR signal transduction^42^) of sEGFR was detected by mass photometry (SI Appendix Fig. S16). This contrasts with the recent speculation that Cu^2+^ may drive dimerization of EGFR to activate signal transduction^43^. Having excluded copper-stimulated EGFR dimerization as a potential activation mechanism of EGFR signal transduction, we explored a different possibility that fluctuations in extracellular copper levels could lead to Cu^1+^ import and inactivation of EGFR phosphatases, thereby effectively inducing EGFR activation.

The highly homologous (99% catalytic site similarity^44^) proteins PTPN1 and PTPN2 (a.k.a. PTP1B and TCPTP, respectively) are EGFR pY1068 (and probably Y1086^45^) phosphatases^46–48^. Previous studies assessing ex vivo^49,50^ and in vitro (cellular)^43,51^ environmental exposure to aberrant levels of copper have suggested metallo-inactivation of PTPN1, via an unknown molecular mechanism. Relatedly, it was previously shown that PTPN1 overexpression was sufficient to partially block EGFR phosphorylation induced by cell membrane-permeable copper-complexes in glial cells^27^. These observations prompted us to consider PTPN1 and PTPN2 as alternative candidates for the copper-sensor(s) responsible for relaying changes in intracellular copper levels into EGFR signaling responses. We subsequently screened siRNA knockdown of several PTPs – including established EGFR phosphatases PTPN1, PTPN2, PTPRJ and PTPRG^46^ – and assessed copper-stimulated EGFR activation by measuring *EGR1* expression level changes. *PTPN2* knockdown significantly diminished copper-stimulated *EGR1* activation in A549 cells; the only PTP knockdown to do so (Fig. 4A). This result supports a mechanism of copper-driven activation of the EGFR signaling via PTPN2 inactivation in A549 cells.

**Figure 4.**
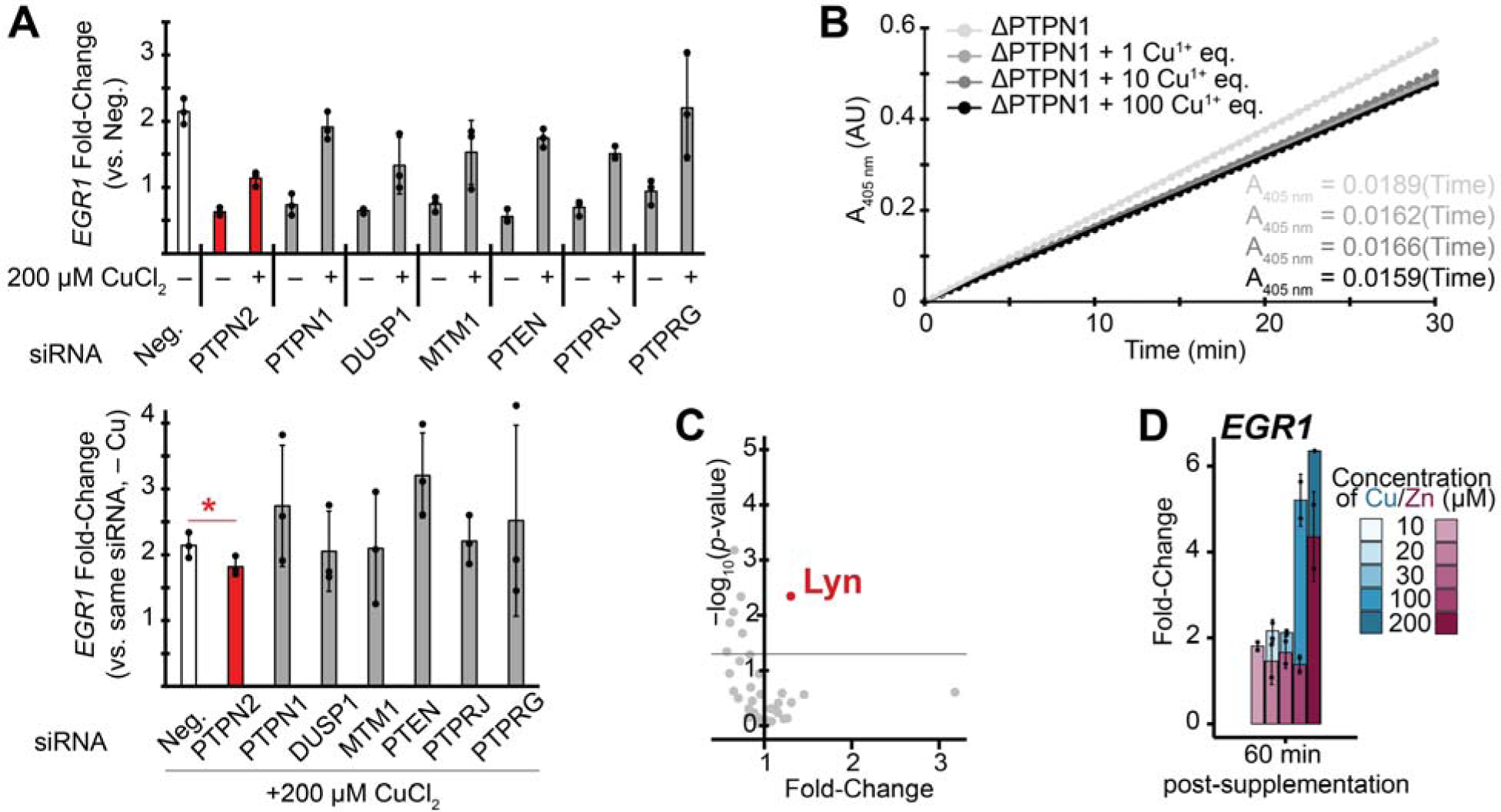
Inhibition of PTPN2 is associated with copper-stimulated activation of EGFR. (*A*) Evaluation of the effects on *EGR1* expression levels of 200 μM CuCl_2_ (60 minute treatment) following siRNA knockdown of various PTPs in A549 cells. Depicted are two interpretations of the results. (*top*) All fold-changes are reported relative to the negative control siRNA transfect without CuCl_2_ supplementation. (*bottom*) Each result is reported as the fold-change between the denoted siRNA knockdown after 60 min 200 μM CuCl_2_ supplementation and the siRNA knockdown by itself. (*n* = 3 replicates per condition). (*B*) *p*NPP (*p*-nitrophenyl phosphate) assay of ΔPTPN1 PTP activity following addition of various amounts of Cu^1+^ (*n* = 12 replicates per condition). (*C*) Proteome profiler antibody array volcano plot depicting statistically-significant increases in phosphorylation levels following HEK 293T supplementation with 20 μM ZnCl_2_. (*n* = 2 replicates for each condition). (*D*) Changes in A549 *EGR1* expression levels 60 minutes after ZnCl_2_ supplementation. The fold-changes are overlayed on the fold-changes induced by CuCl_2_ supplementation for reference. (*n* = 2 replicates per condition).

PTPN1 and PTPN2 dephosphorylate signal transduction proteins to attenuate signaling via a highly-acidic active site cysteine. While Cu^2+^ and other divalent metals are known to inactivate PTPN1, to the best of our knowledge, no biochemical assessment of Cu^1+^-binding nor -inactivation of PTPN1 has been performed^52–54^. As the PTPN1 and PTPN2 catalytic domains are highly homologous, and considerably more metallo-inactivation experiments have been performed on PTPN1, we recombinantly expressed and purified the PTPN1 catalytic domain (ΔPTPN1; residues 1-301), and subsequently evaluated Cu^1+^-binding. We found that addition of excess Cu^1+^ to ΔPTPN1 resulted in the protein binding 5.4 ± 0.47 (*n* = 3) equivalents, as measured by ICP-MS, and ΔPTPN1 activity was inhibited following incubation with Cu^1+^ (as assessed by the *p*-nitrophenyl phosphate (PNPP) assay, Fig. 4B). The active site cysteine mutant C215S ΔPTPN1 bound 4.6 ± 0.26 (*n* = 3) Cu^1+^ equivalents, strongly indicating that the active site cysteine residue binds Cu^1+^, and that the mechanism of the Cu^1+^-driven inhibition involves Cu^1+^ binding to the active site cysteine. By contrast, mutation of C121S (a cysteine residue not directly involved in enzyme catalysis^55^) induced no change in Cu^1+^ binding stoichiometry (5.6 ± 0.14, *n* = 3). We therefore confirmed that copper ions can biochemically modulate PTPN1/2 activity via binding to the catalytic active site cysteine.

Finally, we sought to evaluate specificity of copper-driven PTP inhibition, first relative to the chemically-similar bioavailable metal zinc. Similar to past studies, we detected no significant increase in CREB nor EGFR phosphorylation following addition of the same concentration of ZnCl_2_ as added for CuCl_2_ (20 µM) (Fig. 4C, SI Appendix Fig. S17). Moreover, we found ZnCl_2_ supplementation drove weaker activation of *EGR1* expression (and therefore, weaker EGFR activation) than CuCl_2_ in A549 cells (Fig. 4D). Indeed, previous studies of multiple cell lines found that while cell membrane-permeable Cu^2+^-complexes induced strong EGFR activation, the Zn^2+^-complexes (or ZnCl_2_ supplementation) induced little to no effect^27,56^. Thus, PTP inactivation is more sensitive to copper level fluctuations than zinc, particularly in the physiological range of concentrations.

We then evaluated specificity of copper-driven PTP inhibition relative to *general* PTP inhibition. We compared the CuCl_2_ proteome profiler antibody array results to signal transduction activation following addition of 20 µM pervanadate solution, which serves as a general PTP inhibitor (via oxidation of PTP active site cysteines). While copper supplementation drove significant repression of AKT and p70S6K phosphorylation, presumably indicating repression of mTOR signaling, pervanadate (perhaps unsurprisingly) drove activation of virtually all signaling proteins evaluated (SI Appendix Fig. S18). Thus, the copper signaling responses elicited by PTPN2-inactivation in A549 cells do not reflect a broad PTP inhibition mechanism, but rather drive more specific signaling activation.

## Discussion

Toxicological studies assessing ex vivo^49,50^ and in vitro (cellular)^43,51^ environmental exposure to aberrant levels of copper proposed metallo-inactivation of the ubiquitous, highly-expressed PTPN1 via an unknown molecular mechanism. The previous literature offers conflicting reports on whether or not low-level copper concentration fluctuations – such as occur under normal, physiological conditions – affect this pathway^27,43,56^. Relatedly, our findings represent two breakthroughs.

First, we have identified PTPN2 as a mammalian copper receptor capable of relaying changes in physiological copper levels into signal transduction responses. Regarding the apparent copper-inactivation specificity of PTPN2 over PTPN1, a more comprehensive analysis of PTP siRNA knockdowns in MCF7 breast cancer cells also found PTPN2 knockdown significantly diminished EGFR activation by EGF, while PTPN1 knockdown had no effect^47^. Cell imaging experiments found PTPN1 localized almost exclusively to the perinuclear region, in contrast to PTPN2, which was more diffuse throughout the cell^47^. Consistently, we propose the imported copper is sensed by PTPN2 due to its localization near the cell membrane. We therefore assign PTPN2 as the physiological sensor of copper but note that due to the highly homologous nature of PTPN1 and PTPN2, we do not exclude the possibility that PTPN1 can also function in such a role in different cell contexts.

Second, we have established a link between CREB activation and *CTR1* transcriptional suppression (Fig. 5). Given the superphysiological copper supplementation levels required to drive significant *CTR1* transcriptional repression, it seems implausible that this mechanism exists purely to regulate subtle fluctuations in physiological copper levels, and consequently, our results do not contradict the current mechanistic model that the dominant mechanism of copper-stimulated CTR1 expression regulation occurs at the post-translational level.^15^ However, processes such as neuronal differentiation (as modeled by PC12 cells treated with nerve growth factor, NGF) involve strong, sustained CREB activation^57^, substantial intracellular copper accumulation^6,20^, and strong *CTR1* transcriptional repression (via an unknown regulatory mechanism)^6^. Thus, we propose CREB activation may be the underlying driver of this *CTR1* repression. Interestingly, both the newly discovered pathway reported here and copper-driven inhibition of PDE3B (which leads to elevated cAMP levels, as known to drive CREB activation) converge on cAMP and CREB regulation to connect copper availability with cell growth, proliferation, and differentiation.

**Figure 5.**
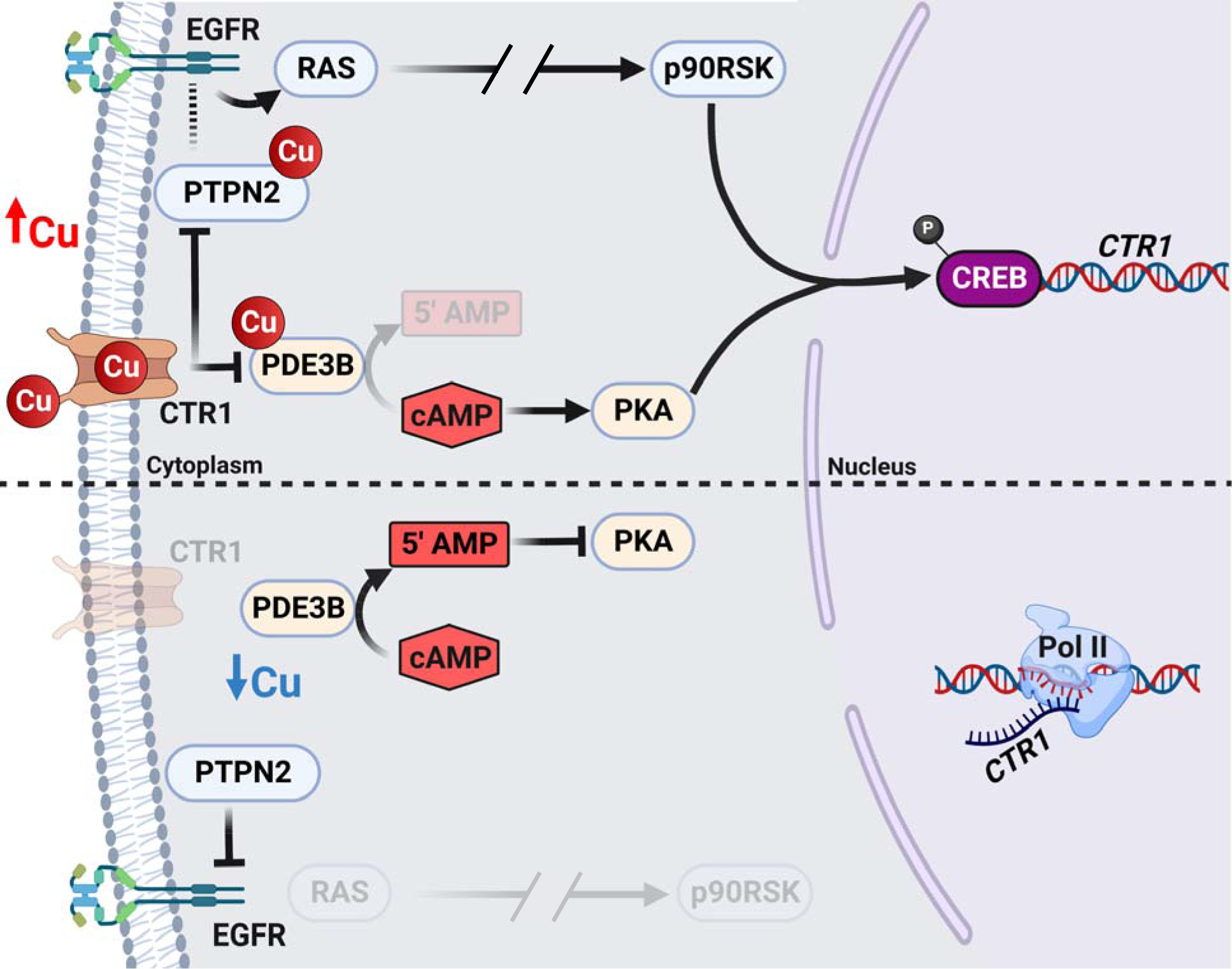
Schematic representation of copper signaling and *CTR1* repression. (***top***) Proposed mechanism of copper-stimulated EGFR activation, with downstream CREB TF activity and transcriptional repression of *CTR1* to regulate intracellular copper levels and copper-stimulated cAMP signaling (***middle*** and ***bottom***). Figure created with BioRender.com.

Finally, EGFR, CREB, and serum/tissue copper depletion are all targets in anti-cancer treatments. Our discovery of a signaling axis involved in *CTR1* repression (a known cause of platinum anticancer drug resistance) represents discovery of a potential targetable axis for synergistic drug efficacy.

## Methods

Unless otherwise noted, all materials were obtained from ThermoFisher. Cetuximab antibodies were a kind gift from Dr. Bruce Marc Bissonnette. CREB Inhibitor 666-15 was purchased from Sigma-Aldrich and dissolved in DMSO to establish a 17.14 mM stock. All volcano plots were generated in RStudio. For statistical-significance reported on images, standard asterisk notations apply, where one asterisk denotes *p*-value < 0.05, two asterisks denote *p*-value < 0.01, and three asterisks denote *p*-value < 0.001. The term “biological replicate” indicates discrete, separate samples; the same sample was never measured repeatedly.

### Cell Culture

A549 and HEK 293T cells were cultured at 37°C, 5% CO_2_, 90% humidity in CellXpert C170i Cell Culture Incubators. Culture media was Dulbecco’s Modified Eagle Medium (DMEM) supplemented with 10% Fetal Bovine Serum (FBS) and 1% Penicillin-Streptomycin (10,000 U/mL) (this combined solution is referred to as supplemented-DMEM, sDMEM). Sub-culturing was performed by aspirating the media, washing with an equivalent volume of Dulbecco’s phosphate-buffered saline (DPBS), aspirating the DPBS, washing with one-fifth the original media volume 0.25% Trypsin-EDTA, aspirating, incubating for one minute in the cell culture incubator, and then resuspending in sDMEM. Cells were seeded at a 1:4 ratio of cell resuspension solution to fresh sDMEM, and were maintained under 95% confluency. Total sDMEM volume used for 35 mm (6 well plate), 100 mm, and 150 mm Corning cell culture plates was 2 mL (per well), 10 mL, and 20 mL, respectively. Cells were verified mycoplasma negative using the MycoProbe Mycoplasma Detection Kit (R&D Systems). For serum-starvation experiments, cell culture sDMEM was replaced with DMEM for times indicated in figures, and cells were subsequently either treated with pre-warmed (to 37°C) DMEM or DMEM supplemented with CuCl_2_ at the stated concentrations. For gefitinib/cetuximab treatments, cells were pre-treated with either 2 µM gefitinib in sDMEM for 3 hours or 80 μg/mL Cetuximab in sDMEM for 4 hours in the cell culture incubator. For subsequent Cu-treatment of EGFR-inhibitor pretreated cells: Cu-supplemented, EGFR-inhibitor supplemented (same concentrations) sDMEM was pre-warmed (to 37°C) and subsequently used to replace the cell supernatant media.

MDA-MB-468 cells were cultured by Dr. Olufunmilayo Olopade’s group in RPMI-1640 supplemented with 10% FBS and antibiotic-antimycotic solution (Gibco). Cells were handled and treated (with TM- and/or DMSO) as described elsewhere^58^.

### Pervanadate Generation

To generate the pervanadate solution for PTP inhibition, we first generated a fresh stock of 100 mM sodium orthovanadate in water. 20 μL of this solution was mixed with 23 μL 30% H_2_O_2_ and 157 μL of DMEM, which was then incubated at room temperature (in the dark) for 30-45 minutes to prepare a 10 mM pervanadate stock. Pervanadate solutions were generated immediately prior to cell treatment.

### Antibody Array and Associated Imaging

Cells were seeded in 150 mm plates and treated at ∼80% confluency. Following serum-starvation, supernatant media was removed and replaced with either DMEM or metal-supplemented DMEM. The plates were subsequently put back into the cell culture incubator for the times noted in the text. Cells were harvested, lysed, and lysates assessed with the Proteome Profiler Human Phospho-Kinase Array Kit (R&D Systems) and C-Series Human EGFR Phosphorylation Antibody Array 1 Kit (RayBiotech) according to manufacturer protocols. Imaging was conducted with the FluorChem R System; experimental and control membranes were imaged for identical exposure durations to facilitate normalization of chemiluminescence intensities across membranes. Integrated pixel densities were measured in ImageJ; pixel densities were subsequently background corrected and normalized to reference control spot-antibody pixel density intensities.

### ChIP-seq Data Analysis

ChIP-seq BigWig files were accessed from the ChIP-Atlas database. Wiggletools v1.2.11 was used to calculate the mean of BigWig files as indicated in the text, which was output as a Wig file. Wig files were subsequently converted to BigWig files using the ucsc-wigtobigwig package (v377). BigWig files were visualized using IGV.

### KAS-seq and Data Analysis

The KAS-seq labeling and isolation procedure was followed as previously described^59^, with the following changes. Cell culture media was removed and subsequently replaced with CuCl_2_-supplemented (or CuCl_2_-unsupplemented control) sDMEM for times indicated in the volcano plots. Following incubation, 500 mM N_3_-kethoxal (in DMSO) stock was added directly to the cell media to a final concentration of 5 mM. After addition, the solution was vigorously shaken to promote solvation of the moderately-soluble N_3_-kethoxal. Dual index library construction used an Accel-NGS Methyl-Seq DNA library kit; libraries were sequenced at the Genomics Facility (University of Chicago) via single-end Illumina NovaSeq 6000 (SP flowcell, 100 bp cassette) sequencing. Reads from two sequencing runs were catenated, and then processed following the KAS-pipe data processing pipeline^59^ (mapped to human genome build hg19) on a Lenovo Thinkstation P920 workstation.

### RNA extraction and RNA-qPCR

Total RNA was isolated using either TRIzol according to provided instructions, or a combination of TRIzol and the RNA Clean & Concentrator-5 (RCC-5) kit (Zymo Research), according to the Zymo instructions for combination Trizol/Zymo-Spin-Column RNA isolation. Samples prepared using the RCC-5 kit were also treated with in-column DNase I following Zymo instructions. cDNA synthesis was performed via either Maxima First Strand cDNA Synthesis Kit for RT-qPCR (with dsDNAse added), or High Capacity cDNA Reverse Transcription Kit (Applied Biosciences). RNA levels were quantified by determining cycle threshold (Ct) values of corresponding cDNAs with the QuantStudio 6 Pro system (Applied Biosciences, A43180) and FastStart Essential DNA Green Master (Roche) at a final concentration of 1X, with ∼500 nM forward and reverse primers, in a final reaction volume of 20 μL per well in MicroAmp Optical 96-Well Reaction Plates (primer sequences provided in SI Appendix Table S6). Samples were amplified with the following run method: 10 min hold at 95°C, 40x cycles of 20 sec at 95°C, 20 sec at 60°C, and 20 sec at 72°C (during which fluorescence measurements occurred), before a final 15 sec at 95°C and 1 min at 60°C. The cycle was completed with a melting curve while measuring fluorescence. All biological replicates comprised at least 2 technical replicates. All transcripts were processed using the 2^−ΔΔCt^ relative expression normalization method, using *GAPDH* as an internal reference standard. Copper treatment RNA-qPCR experiments compared metal-treated to control sDMEM-treated values. siRNA treatment experiments compared cDNA generated following experimental siRNA treatment to cDNA generated from Silencer™ Select Negative Control No. 1 siRNA-transfected cells. Gefitinib pretreatment experiments compared gefitinib-pretreated, Cu-treated to gefitinib-pretreated cell cDNA. Two-tailed, paired T-test *p*-values were calculated in RStudio, using the following packages: dplyr v1.0.7, tidyverse v1.3.1, readxl v1.3.1, ggprism v1.0.3. Example code is provided in the Supplementary Materials.

### CREB1 and PTP Knockdown

Knockdown of CREB1 and PTPs was performed via transfection of Silencer Select siRNAs using Lipofectamine™ RNAiMAX Transfection Reagent. Specifically, cells were seeded in sDMEM lacking antibiotics ∼24 hours prior to transfection to generate 6 well plates in which individual wells were ∼40-50% confluent at time of treatment. Per cell treatment: 9 μL of Lipofectamine was diluted into 150 μL of Opti-MEM at room temperature, and separately, 3 μL of 10 μM siRNA was diluted into 150 μL of Opti-MEM. The diluted siRNA was then mixed with the diluted Lipofectamine in a 1:1 ratio, and incubated at room temperature for 5 minutes. 250 μL of the Lipofectamine/siRNA mixture was added to the 2 mL of cell media in the well, and then quickly mixed back and forth to promote homogenization. Cells were then put back into the cell incubator for 24 hours prior to treatment and/or RNA extraction and RNA-qPCR measurement.

### Constitutive CREB Expression Vector Transfection

The CREB Dominant-Negative Vector Set was purchased from Takara Bio, and the vectors subsequently transformed into DH5α Mix and Go! competent *E. coli* (Zymo Research) and plated onto LB-agar plates with 50 μg/mL kanamycin and colonies grown overnight at 37°C. Vector transformed colonies were picked and grown in a 10 mL LB culture with 50 μg/mL kanamycin (37°C with vigorous shaking). This culture was used to inoculate overnight 1 L cultures of LB with 50 μg/mL kanamycin (grown at 37°C with vigorous shaking). The plasmid was subsequently purified using a QIAGEN plasmid maxi kit following manufacturer protocol.

A549 cells in antibiotic-free sDMEM were transfected with 2.5 μg of purified pCMV-CREB Vector (corresponding to a plasmid for constitutive CREB expression) using a Lipofectamine 3000 reagent kit, following the manufacturer protocol. As a negative control, cells were transfected (following the same procedure) with the DsRed-EGFP in a pcDNA3.1 HisC vector^60^.

### MDA-MB-468 RNA-seq and Data Analysis

Total RNA was extracted from MDA-MB-468 cells via Trizol extraction; the RiboMinus Eukaryote Kit was then used according to manufacturer protocol to remove ribosomal RNA. RNA were then used for library construction using a SMARTer Stranded RNA-seq kit (Takara, 634839) according to the manufacturer protocol. Paired-end high-throughput sequencing was subsequently performed at the University of Chicago Genomics Facility on an Illumina HiSeq 4000 sequencer.

Reads were trimmed using the KAS-pipe paired-end trim_adapter.sh script. Trimmed reads were aligned to STAR reference genomes (using STAR version 2.7.9a) generated from the human genome build hg38. Aligned reads were then processed to remove (*i*) rows “N_ambig”, “N_multima”, and “N_noFeat” and (*ii*) transcripts with 2 or fewer average read counts. Unstranded column reads were then processed for differential expression analysis via DESeq2 v1.36.0. Pathway/process enrichment analysis of significantly differentially-expressed transcripts was performed via Metascape^61^, while GSEA was performed using GSEA v4.2.3.

### A549 RNA-seq and Data Analysis

Total RNA was isolated using a combination of TRIzol and the RCC-5 kit (Zymo Research), according to Zymo’s instructions for combination Trizol/Zymo-Spin-Column RNA isolation. Samples prepared using the RCC-5 kit were also treated with in-column DNase I according to provided Zymo instructions. Total isolated RNA was subjected to two rounds of poly(A)-enrichment using the Dynabeads mRNA DIRECT Kit according to the manufacturer’s instructions twice on each sample.

Library preparation was performed with 5 µL of twice poly(A)-enriched sample using the NEBNext Ultra II Directional RNA Library Prep Kit for Illumina (New England BioLabs) according to manufacturer’s instructions for “Protocol for use with Purified mRNA or rRNA Depleted RNA”. Paired-end high throughput sequencing was subsequently performed on an Illumina NextSeq 550. FastQ files from each sample across runs were subsequently combined. Adapter trimming and basic QC was carried out with Trim Galore. Alignment to the human hg38 reference genome was carried out with HISAT2. Duplicate read removal was carried out with Samtools. Transcript identification and read counting was performed with featureCounts according to gene transfer format files provided by release 107 of the Ensembl database. Transcripts with fewer than 0.74 average reads per replicate across all samples were excluded from analysis. Differential expression analysis was performed by the DESeq2 R package DESeq2 v1.34.0. Pathway/process enrichment analysis of significantly differentially-expressed transcripts was performed via Metascape^61^.

### TCGA and GTEx RNA-seq Data Analysis

Lung adenocarcinoma (LUAD) TCGA data was accessed via the Xenabrowser server^62^ – transcript read counts were converted from “log_2_(norm count + 1)” to “norm count + 1” prior to statistical analysis. *p*-values were generated in Excel via the t-Test: Two-Sample Assuming Unequal Variances function.

GTEx Analysis V8 release gene read counts were downloaded from the GTEx Portal. Read counts were first separated by tissue into discrete datasets, which were subsequently processed and analyzed separately. Next, transcripts with 2 or fewer average read counts were removed. Next, differential expression analysis (comparing tissue samples against an arbitrarily chosen sample from the same tissue) via DESeq2 v1.36.0 was performed to generate normalized read counts. Transcript correlations and plots (as well as associated statistical analysis) were subsequently generated in RStudio from the DESeq2 normalized read counts using ggpubr v0.4.0. Example code is provided in the Supplementary Materials.

### Cloning and Expression of sEGFR

Recombinant human EGFR extracellular domain (residues 1-642) was expressed in insect cells as secreted protein using an approach described previously^63^. Briefly, human EGFR extracellular domain (residues 1-642) followed by an HRV3C cleavage site and a C-terminal 6x-histidine tag was subcloned into a pVL1393 vector. To generate the baculovirus, the constructed plasmid was co-transfected with the linearized baculovirus DNA (Expression Systems, 91-002) using Cellfectin II (Thermo Fisher, 10362100). Baculovirus was subsequently amplified in Sf9 cells (Thermo Fisher, 12659017) cultured in Sf-900 III SFM medium (Gibco, 12658019), supplemented with 10% heat-inactivated fetal bovine serum (Cityva, SH30396), 2 mM L-Glutamine (Cityva, SH3003402), and 10□μg/mL gentamicin at 27°C. Large-scale protein expression was performed by infection of High-Five cells (Thermo Fisher, B85502) in Insect-XPRESS medium (Lonza, BELN12-730Q) medium at a cell density of 2.0□×□10^6^ cells/ml for 64 to 70□h at 27□°C.

### Purification of sEGFR

The EGFR signal peptide (residues 1-24) was cleaved after secretion. The medium containing secreted EGFR (25-642) was collected and centrifuged at 900x*g* for 15□min at room temperature. The supernatant was transferred into a beaker and mixed with buffer and salt reagents; the final mixture contained 50□mM Tris, pH 8.0, 5□mM CaCl_2_ and 1□mM NiCl_2_. The mixture was stirred at 130 rpm for 30□min at room temperature. After a centrifugation (8000x*g*, 30□min), the cleared supernatant was incubated with Ni-NTA resin (Thermo Scientific, 88223) for 4 hours at room temperature. The resin was collected using a glass Buchner funnel and rinsed with HBS buffer (10 mM HEPES, pH 7.2, 150 mM NaCl) containing 10□mM imidazole, before transferring to a gravity column (Bio-Rad, 7372512). The protein was eluted with HBS buffer containing 200□mM imidazole. The eluent was concentrated and further purified by size-exclusion chromatography (Superdex 200 10/300 GL; GE Healthcare) equilibrated in a buffer containing 20□mM HEPES, pH 7.5 and 100□mM NaCl. Cleavage of the His-tag by HRV3C protease (Takara, 7360) followed the manufacturer’s protocol. HRV3C protease and cleaved His-tag were removed from sEGFR by a second size-exclusion chromatography (Superdex 200 10/300 GL) run. The successful removal of His-tag was validated by western blot using the anti-His-tag antibody (Genscript, A00186-100).

### Phen Green Competition Experiments

For fluorescence measurements, 1 mg aliquots of Phen Green were dissolved in 472 μL of DMSO – from this stock, a 3 μM Phen Green stock in 20 mM HEPES, 100 mM NaCl, pH 7.5 was prepared. Invitrogen 96-well Fluorescence-based Assay Microplate wells were filled with 600 μL of either (*i*) 20 mM HEPES, 100 mM NaCl, pH 7.5 buffer control, (*ii*) 3.64 μM sEGFR, 1 μM Phen Green (dissolved in DMSO) in 20 mM HEPES, 100 mM NaCl, pH 7.5, and (*iii*) 3.64 μM sEGFR, 1 μM Phen Green in DMSO in 20 mM HEPES, 100 mM NaCl, pH 7.5 with a screen of CuCl_2_ concentrations (denoted in the respective figures) ranging from 0 equivalents (relative to Phen Green) to 1 equivalent. Fluorescence measurements were performed on a Synergy HTX Multi-Mode Reader, using an excitation wavelength of 485 nm and an emission wavelength of 528 nm (with the optics in the top position). Following measurement, to all wells 3 μL of 5 μM CuCl_2_ in 20 mM HEPES, 100 mM NaCl, pH 7.5 were added; samples were then incubated at room temperature for 15 minutes prior to subsequent fluorescence measurement. Δ_Fluorescence_ titration curves were then generated by averaging 5 measurements for each CuCl_2_ titration concentration, and defining the 0 μM CuCl_2_ added measurement as “0” and the lowest fluorescence intensity measurement as “1”. Curve fitting was performed using the Matlab Curve Fitting App to the equation 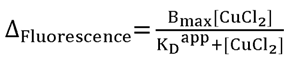, where [CuCl_2_] corresponds to the added concentration of CuCl_2_ in 20 mM HEPES, 100 mM NaCl, pH 7.5, B_max_ corresponds to the number of Cu binding sites on the protein, and K_D_^app^ is the apparent Phen Green K_D_ (corresponding to the true Phen Green KD when [sEGFR] = 0, assuming negligible Cu^2+^ binding by the buffer) This value was used to determine the sEGFR Cu^2+^ as described below.

We first assumed that there was no “free” solution Cu^2+^: all Cu^2+^ added to the solution was bound by sEGFR or Phen Green. Then, from the equation 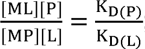; where [ML], [P], [MP], and [L] correspond to the Phen Green-bound Cu^2+^, apo-sEGFR, sEGFR-bound Cu^2+^, and apo-Phen Green concentrations, respectively, while K_D(P)_ and K_D(L)_ are the Cu^2+^ K_D_ values for sEGFR and Phen Green, respectively; [L] = [ML] when [MP] = K_D_^app^ (this is just a substitution of [MP] for “free” solution Cu^2+^, or [M] in this case, into the definition of K_D_). Consequently, when [MP] = K_D_ ^app^, the equation may be simplified to 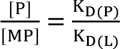 or 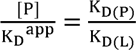. From this equation, we calculated K_D(P)_.

### ICP-MS

Metal-bound protein solutions (sEGFR or ΔPTPN1) were precipitated to release bound metals via addition of 5% HNO_3_. The protein precipitate was subsequently pelleted by centrifugation. Sample measurements were performed via an Agilent 7700x ICP-MS in He mode, and were subsequently analyzed using ICP-MS Mass Hunter version B01.03. Intensities were correlated to ppb concentrations using multi-element ICP-MS standards (Inorganic Ventures).

### EPR Spectroscopy

For EPR measurements, ∼200 μL of ∼100-200 μM sEGFR was aliquoted into Wilmad quartz X-band EPR tubes and flash frozen in liquid nitrogen. X-band continuous wave (CW) EPR measurements were performed at both the University of Chicago using a Bruker Elexsys E500 spectrometer featuring an Oxford ESR 900 X-band cryostat with a Bruker Cold-Edge Stinger.

### Mass Photometry

Mass photometry experiments were conducted at the University of Illinois at Chicago campus, using a Refeyn Two Mass Photometer. sEGFR samples (final concentration of 100 nM) were incubated overnight at 4°C with the indicated concentrations of CuCl_2_. Samples were mixed on the mass photometer in a 1:1 ratio of protein sample to buffer.

### Cloning and Expression of ΔPTPN1 constructs

The ΔPTPN1 and C215S ΔPTPN1 constructs were cloned into a pET-28a(+)-TEV vector, featuring C-terminal TEV protease cleavage site followed by (His)_6_-tag, by Genscript. The C121S ΔPTPN1 mutant was generated from the ΔPTPN1 vector using a QuikChange II Site Directed Mutagenesis kit (Agilent), and the mutant sequence verified by whole plasmid sequencing (Azenta). Vectors were transformed into BL21 Star (DE3) *E. coli* via heat shock and were plated onto Kan50 plates (50 μg/mL kanamycin in LB-agar) for antibiotic selection. Vector-transformed *E. coli* colonies were subsequently picked and grown at 37°C in 10 mL overnight starter LB-cultures with 50 μg/mL kanamycin. After overnight growth, 3 mL of cell suspension was used to inoculate 1 L of LB media + 50 μg/mL kanamycin. This culture was grown with vigorous shaking at 37°C until the culture reached an OD_600_ of ∼0.5, at which point IPTG was added to a final concentration of 200 μM and the cell culture temperature was lowered to 15°C. Following overnight protein expression, the cells were pelleted by centrifugation (4°C, 5000x*g*, 10 min) and the supernatant poured off. Cell pellets were subsequently frozen in liquid nitrogen, and stored at −80°C.

∼10 g cell pellets were thawed and resuspended in 50 mL pre-chilled cell lysis buffer (100 mM Tris-HCl pH 7.5, 100 mM NaCl, 10 mM BME, 5% glycerol) and lysed via sonication. After sonication, the cell solution was moved to a 50 mL conical tube and the cell remains were pelleted by centrifugation (4°C, 30 min, 10000x*g*). The supernatant was moved to a new 50 mL conical tube and again cleared by centrifugation (4°C, 30 min, 10000xg). The supernatant was moved to a new 50 mL conical tube and held on ice.

2 mL of HisPur Ni-NTA Resin per 10 g cell pellet were washed repeatedly with column buffer (100 mM Tris-HCl pH 7.5, 100 mM NaCl, 10 mM BME, 10 mM imidazole) and resuspended to a final volume of 2 mL with column buffer. The resin was then added to the cleared cell supernatant in lysis buffer, and the supernatant solution was subsequently gently mixed at room temperature for 30 minutes. After 30 min, removed supernatant solution and washed 5x with column buffer (beads were centrifuged for 2 min, 4°C, 700x*g*, supernatant removed, and beads resuspended with column buffer). After the final wash, removed the supernatant solution was removed and 10 mL of elution buffer (100 mM Tris-HCl pH 7.5, 100 mM NaCl, 10 mM BME, 500 mM imidazole) was added. The beads were incubated on ice for 10-15 min with gentle periodic resuspension, and then were centrifuged (2 min, 4°C, 700x*g*). The supernatant solution containing the eluted protein was set aside, and the elution buffer wash and centrifugation repeated. The eluted protein was then stored at 4°C overnight.

The eluted protein was subsequently concentrated using an Amicon 3000 kDa cutoff spin filters via centrifugation (4°C, 4000x*g*). Following concentration, the protein was buffer exchanged into buffer II (100 mM Tris-HCl pH 7.5, 100 mM NaCl, 10 mM TCEP-HCl) using a PD-10 desalting column. To the eluted protein fractions, 25 uL of (His)_6_-tagged TEV-protease (Sigma-Aldrich) was added and the solution incubated overnight at 4°C.

After overnight incubation, ∼1.5 mL of HisPur Ni-NTA Resin per protein elution was washed and resuspended in buffer II. The resin was then added to the TEV-protease cleaved protein solution to remove the TEV-protease as well as the cleaved (His)_6_-tag. The solution was incubated at 4°C with periodic, gentle mixing for 30 minutes, and the beads were then pelleted by centrifugation (4°C, 4000xg). The supernatant containing the purified, (His)_6_-tag-cleaved protein was put into a new tube, and the protein concentration determined by nanodrop (assuming an extinction coefficient of ε_Molar_ = 46410 M^−1^cm^−1^. The protein solutions were moved into 1 mL aliquots in 1.5 mL Eppendorf tubes, which were flash frozen in liquid nitrogen and stored at −80°C.

### ΔPTPN1 Construct Cu^1+^ Binding Assay

To evaluate Cu^1+^ binding by the ΔPTPN1 constructs, the purified, (His)_6_-tag-cleaved protein was thawed on ice and diluted with buffer II to a final concentration of 35 μM. To generate Cu^1+^, we used an adapted method from Padilla-Benavides et al.^64^ Specifically, CuCl_2_ solutions were prepared at the indicated concentrations in buffer II with 100 mM sodium ascorbate included as well, to fully reduce the Cu^2+^ to Cu^1+^. 500 μL of protein solution was mixed with 500 μL of 350 μM CuCl_2_ in buffer II with 100 mM sodium ascorbate; the solution was incubated on ice for 1 hour. After 1 hour, the solution was run over a PD-10 desalting column (held at 4°C) equilibrated with buffer II. The eluted sample was then used to prepare ICP-MS samples.

### ΔPTPN1 *p*-Nitrophenyl Phosphate (PNPP) Assay

To assess PTP activity of the ΔPTPN1 construct, 100 nM purified, His-tag cleaved ΔPTPN1 was aliquoted into a 96 well plate and incubated for 15 minutes with Cu^1+^ solution (generated the same as in the ΔPTPN1 Construct Cu^1+^ Binding Assay) at room temperature. 20 μL of 500 mM PNPP (New England Biolabs) was then added to the 190 μL protein mixture (per well) to generate a final concentration of 47.62 mM PNPP per reaction. This mixture was immediately monitored (with a Synergy HTX Multi-Mode Reader plate reader) for the absorbance at 405 nm, corresponding to the chromogenic product. All readings were normalized to the initial measurement for each replicate.

## Acknowledgments

This work was supported by funding from the National Institutes of Health (R01ES030546 and HG006827) to C.H., R35GM143052 to M.Z., K99ES034084 to M.O.R., the Breast Cancer Research Foundation FP049439 to O.I.O., and University of Chicago Yen Postdoctoral Fellowship to M.O.R. We appreciate the kind support from UChicago genomics facility, Mass Spectrometry Facility, and ARC, especially Dr. C. Jin Qin and Dr. Pieter W. Faber for assisting in the experiments. C.H. is an Investigator of the Howard Hughes Medical Institute. The authors are grateful to Prof. Bruce Marc Bissonnette for the providing Cetuximab antibodies as a kind gift. We thank Prof. Wenbin Lin and Yingjie Fan (University of Chicago) for running the ICP-MS samples.

## Author Contributions

Conceptualization: M. O. R., C. H. Methodology: M. O. R., Y. X., M. Z., C. H. Investigation: M. O. R., Y. X., R. O., C. Y., O. N. P. Z., R. L., T. W., O. K. Funding acquisition: M. O. R., L. L., M. Z., C. H. Supervision: O. I. O., M. Z., C. H. Writing – original draft: M. O. R., Y. X., R. O. Writing – review & editing: M. O.R., C. Y., L. L., M. Z., C. H.

## Supporting Information

### Data availability

High throughput sequencing data are available through the Gene Expression Omnibus (GEO), accession GSE211339 (A549 KAS-seq), GSE210777 (MDA-MB-468 RNA-seq), and GSE214566 (A549 RNA-seq). Metascape gene ontology (GO) enrichment analysis summaries are provided as Tables S4 (MDA-MB-468 TM-treated) and S5 (2 hour, 30 µM CuCl2 treated A549 RNA-seq). DESeq2 differential expression analysis results from the various KAS-seq and RNA-seq experiments are provided as Tables S7 (A549 KAS-seq), S8 (MDA-MB-468 RNA-seq), and S9 (A549 RNA-seq).

### Code availability

Example code and pseudocode used for next generation sequencing data processing is provided in the Supporting Materials section.

### Example qPCR Data Analysis Code

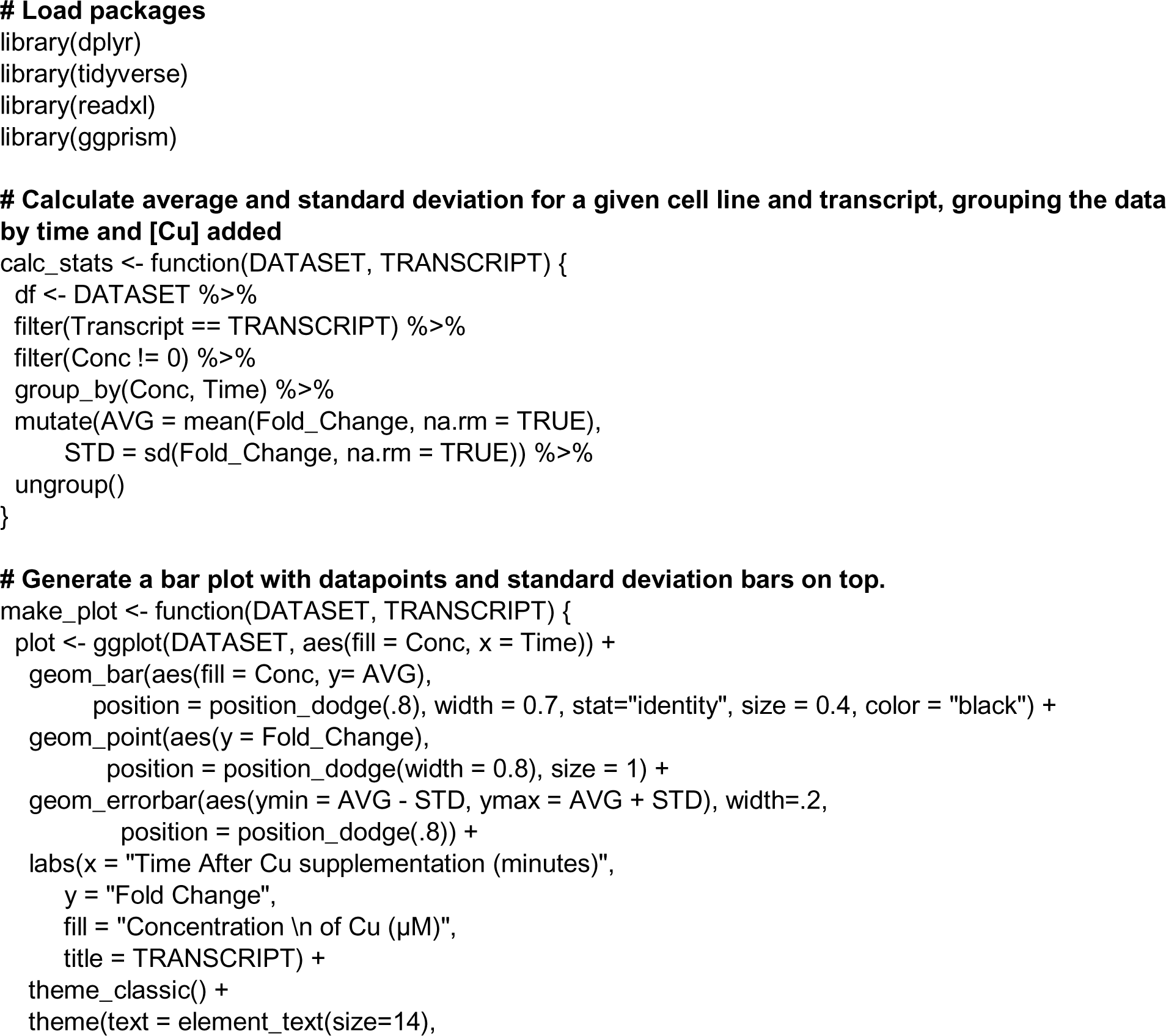
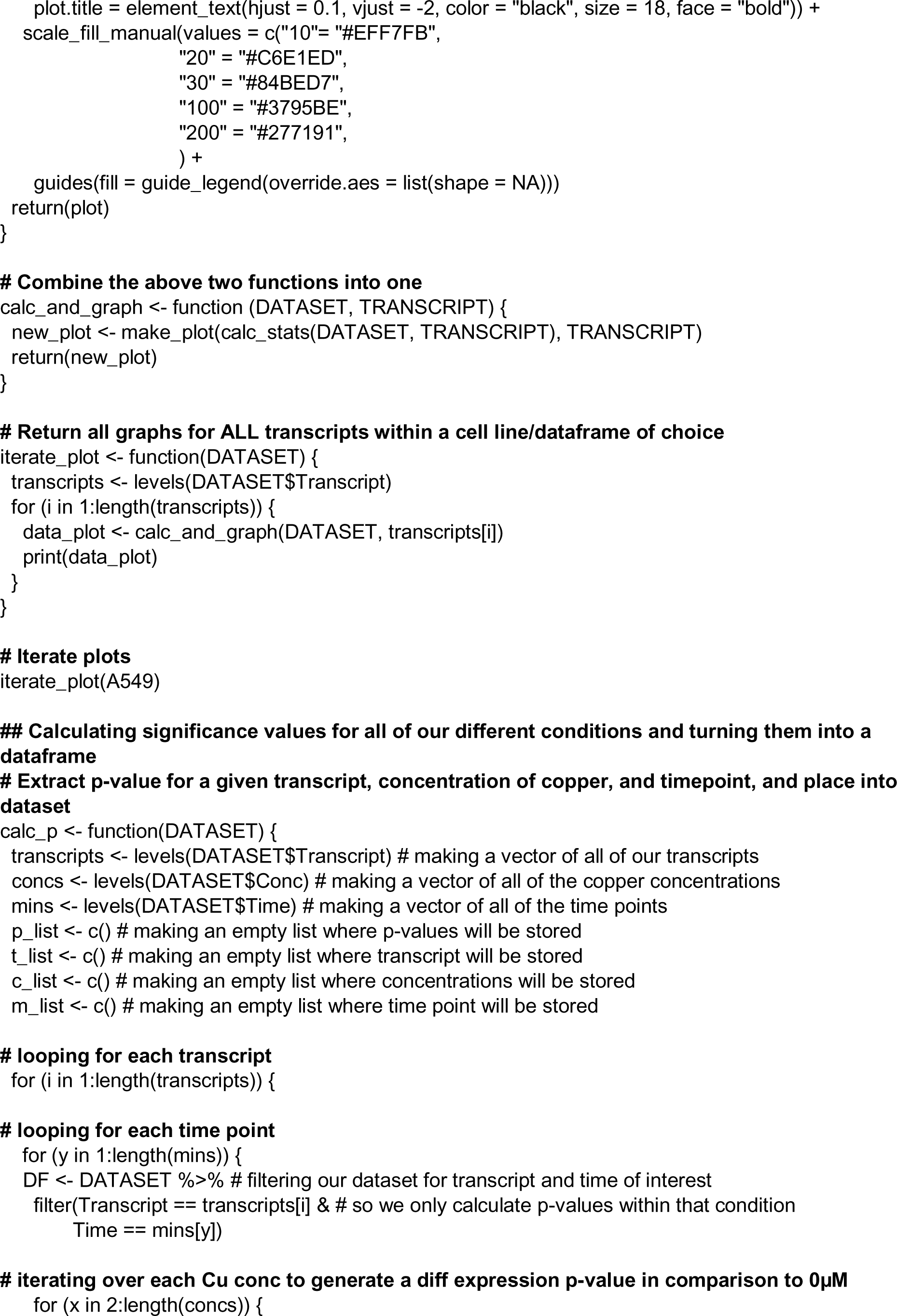
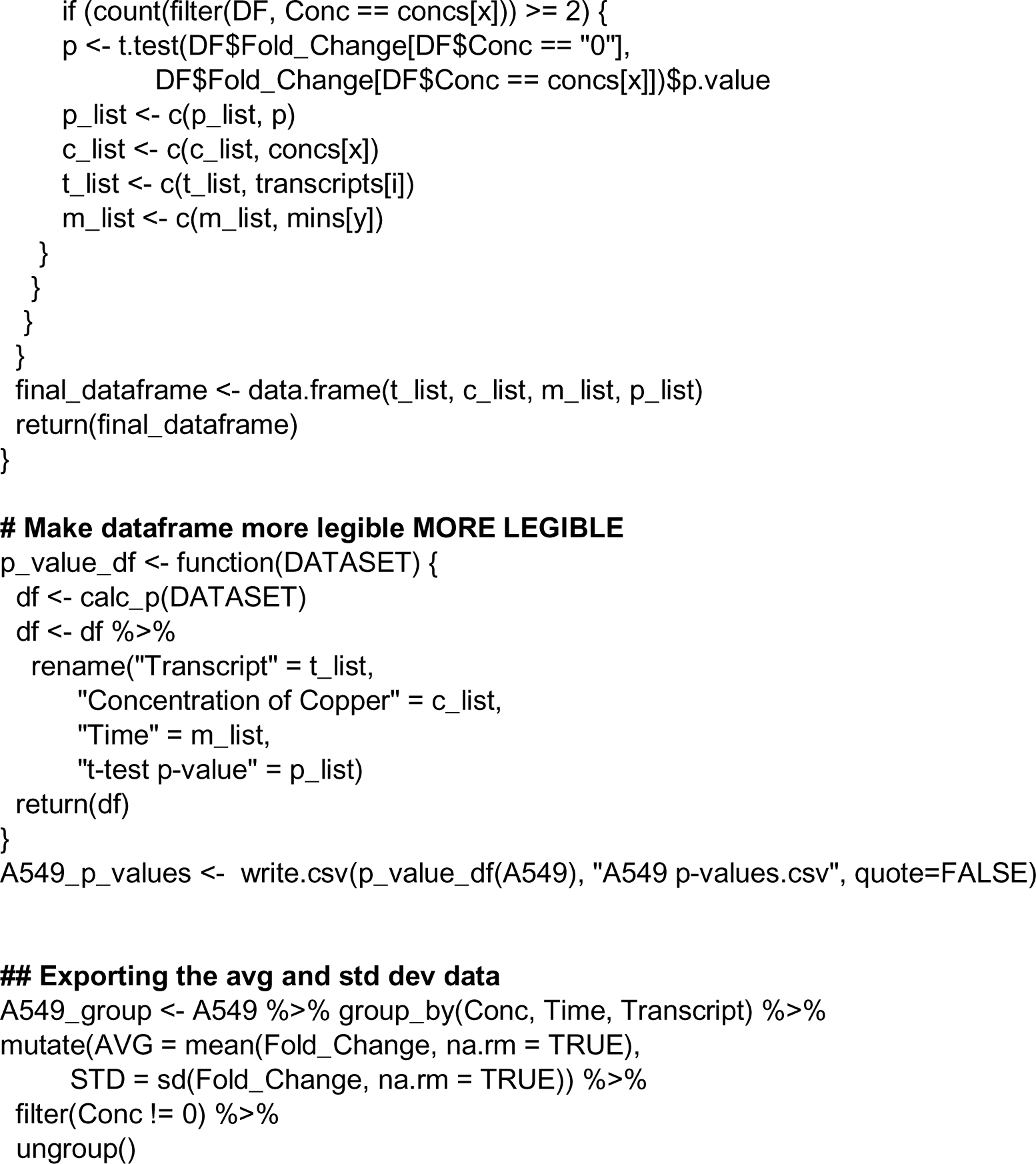

### Example GTEx Data Processing (Psuedo)Code

**#packages**

~~~
library(ggpubr)
library(DESeq2)
~~~

**#read in Annotations**

~~~
sample.df <- read.delim(“GTEx_Analysis_v8_Annotations_SampleAttributesDS.txt”, as.is=TRUE, header=TRUE, row.names=1)
~~~

**#Call RNA-seq samples that pass analysis freeze**

~~~
rnaseq.sample.df <- sample.df[sample.df[’SMAFRZE’]==’RNASEQ’,]
~~~

**#Summary of Tissues**

~~~
as.matrix(sort(table(rnaseq.sample.df[’SMTSD’]), decreasing=TRUE))
~~~

**#Call a specific tissue**

~~~
Tissue.sample.df = rnaseq.sample.df[rnaseq.sample.df[’SMTSD’]==’Tissue - Identifier’,] head(Tissue.sample.df)
~~~

**#Load expression data**

~~~
Counts.df <- read.delim(“GTEx_Analysis_2017-06-05_v8_RNASeQCv1.1.9_gene_reads.gct.gz”,
             as.is=T, row.names=1, check.names=FALSE, skip=2)
gene.names.df <- Counts.df[, ‘Description’, drop=FALSE]
Counts.df <- Counts.df[, !(names(Counts.df) %in% c(’Description’))]
cat(paste(“Number of genes in table:”, dim(Counts.df)[1]))
~~~

**#Call donor IDs from tissue and call tissue specific expression counts**

~~~
transpose.Counts.df <- t(Counts.df)
donor.ids <- rownames(Tissue.sample.df)
Tissue.Counts.df <- transpose.Counts.df[rownames(transpose.Counts.df) %in% donor.ids,]
~~~

**#Export unnormalized counts**

~~~
write.csv(Tissue.Counts.df, “unnormalized.Tissue.counts.csv”)
~~~

**#load count matrix and info table containing samples split into 2 arbitrary conditions**

~~~
dat <- t(read.csv(“unnormalized.Tissue.counts.csv”, header = T, row.names = 1))
info <- read.table(“Tissue_colData.txt”, header = T, sep = ’\t’)
dds <- DESeqDataSetFromMatrix(dat, info, ∼Condition)
~~~

**#remove lowly expressed genes**

~~~
keep <- rowSums(counts(dds)) > “Total number of samples”*2
dds <- dds[keep,]
~~~

**#main DESeq**

~~~
ddsDE <- DESeq(dds)
~~~

**#export normalized read counts**

~~~
normCounts <- counts(ddsDE, normalized = T)
write.csv(normCounts, “normal.Tissue.csv”)
~~~

**#transpose matrix & convert matrix to dataframe**

~~~
transpose_Tissue <- t(Tissue)
Tissue.df <- as.data.frame(transpose_Tissue)
~~~

**#correlate Transcript_1 vs. Transcript_2**

~~~
pdf(file = “Tissue_Transcript_1_vs_Transcript_2.pdf”, width = 10, height = 10)
ggscatter(Tissue.df, x = “ENSG_ID_Transcript_2”, y = “ ENSG_ID_Transcript_1”, alpha = 0.5, col = “blue3”, size = 4,
        xlab = “TRANSCRIPT_2”, ylab = “Transcript_1”, add = “reg.line”, add.params = list(size = 6), conf.int = TRUE,
cor.coef = TRUE,
        cor.method = “pearson”, pch = 20) +
 theme(axis.ticks = element_line(size = 2.5), axis.line = element_line(size = 3.0),
       axis.ticks.length = unit(0.6, “cm”))
dev.off()
~~~

---

**Fig. S1.**
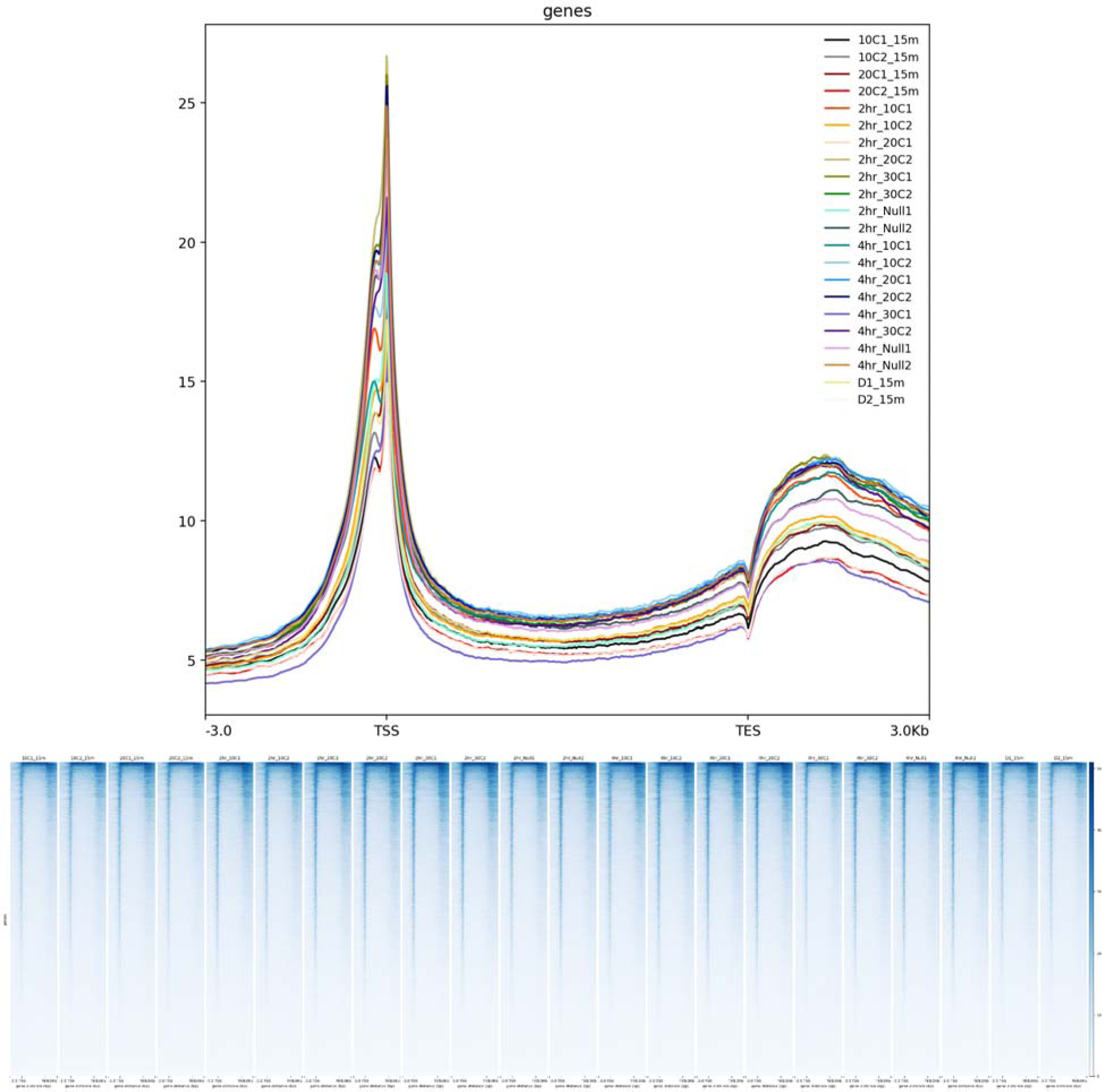
A549 KAS-seq heatmap and profiles. (*top*) Heatmap and (*bottom*) profiles of A549 KAS-seq data, with individual replicates shown separately. The KAS signal shows clear enrichment near the transcription start site (TSS), and to a lesser extent, near the transcription end site (TES). Labels denote the conditions probed for each individual replicate (for example, 10C1_15m was replicate 1 of A549 cells supplemented with 10 μM CuCl_2_ for 15 minutes, 10C2_15m was replicate 2 of the same condition, etc.).

**Fig. S2.**
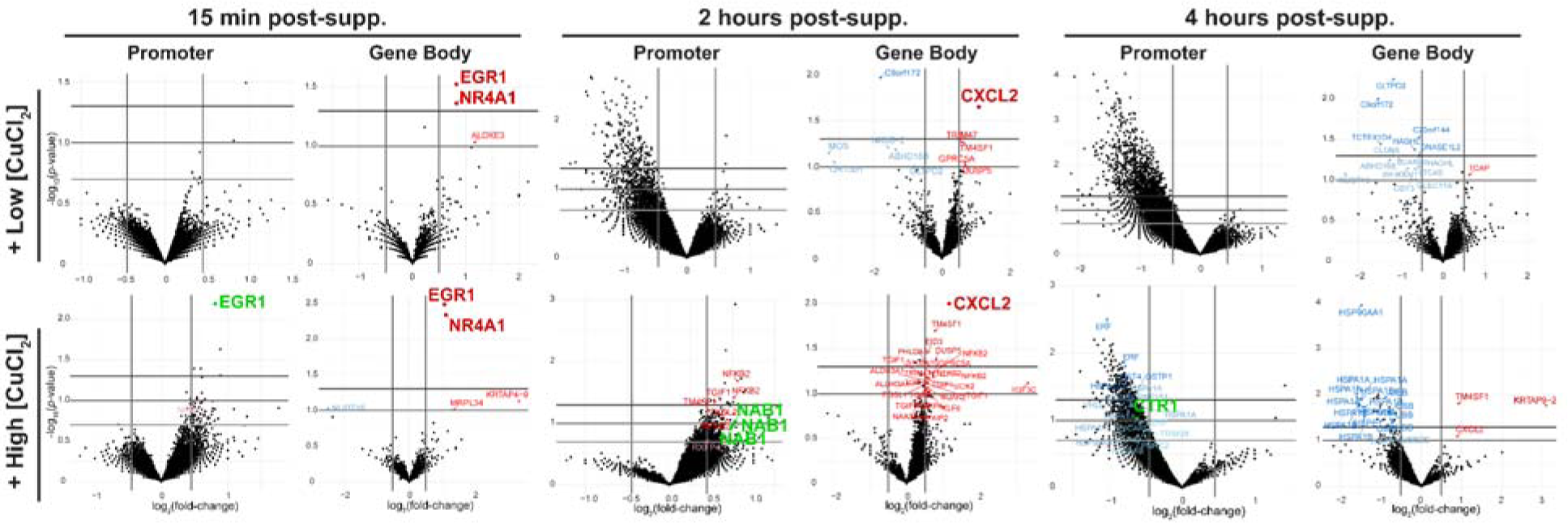
Copper-supplemented A549 KAS-seq volcano plots. Volcano plots compared copper supplemented to unsupplemented cells. Significantly upregulated gene bodies/promoters post-copper supplementation in red, downregulated in blue, and select promoters (*EGR1*, *NAB1* and *CTR1*) in green. *EGR1*, *NR4A1*, *CXCL2, NAB1,* and *CTR1* shown in bold. Labeled promoter data points correspond to up/down-regulated gene bodies at that condition and timepoint. Heavy horizontal lines denote *p*-value = 0.2 (promoter only), 0.1, and 0.05, heavy vertical lines denote |log_2_fold-change| = 0.45 (promoter) or 0.5 (gene body). The multiple *NAB1* promoter datapoints correspond to different promoter region boundaries. For clarity, ubiquitin and heat shock protein differential ssDNA levels omitted from 4-hour high Cu gene body responses (which comprise the unlabeled, significantly repressed data points). In the high Cu datasets, there is excellent agreement for activated/repressed gene bodies with corresponding promoter responses, but not vice versa; promoter KAS-seq measurements are far more sensitive (Supplementary Fig. 1). *n* = 2 biological replicates for each condition.

**Fig. S3.** Copper-stimulated phosphorylation patterns. (*top*) HEK 293T proteome profiler phospho-kinase antibody arrays, (*bottom*) integrated pixel densities reporting chemiluminescence intensity for treatments with or without supplementation of 20 μM CuCl_2_ for 10 minutes (in cell culture incubator: 37°C, 5% CO_2_, 90% humidity). Graph titles indicate the protein name and phosphorylation site (residue) reported. Italicized letters below name of phosphorylation site in graphs corresponds to the location of the antibodies on the membrane array, as indicated by the key in the bottom right. Antibody locations A1,2; A17,18, and G1,2 are reference (control) spots, while G9,10 and G17,18 are negative controls (PBS). Error bars denote standard deviation. *n* = 3 biological replicates for each condition.

**Fig. S4.**
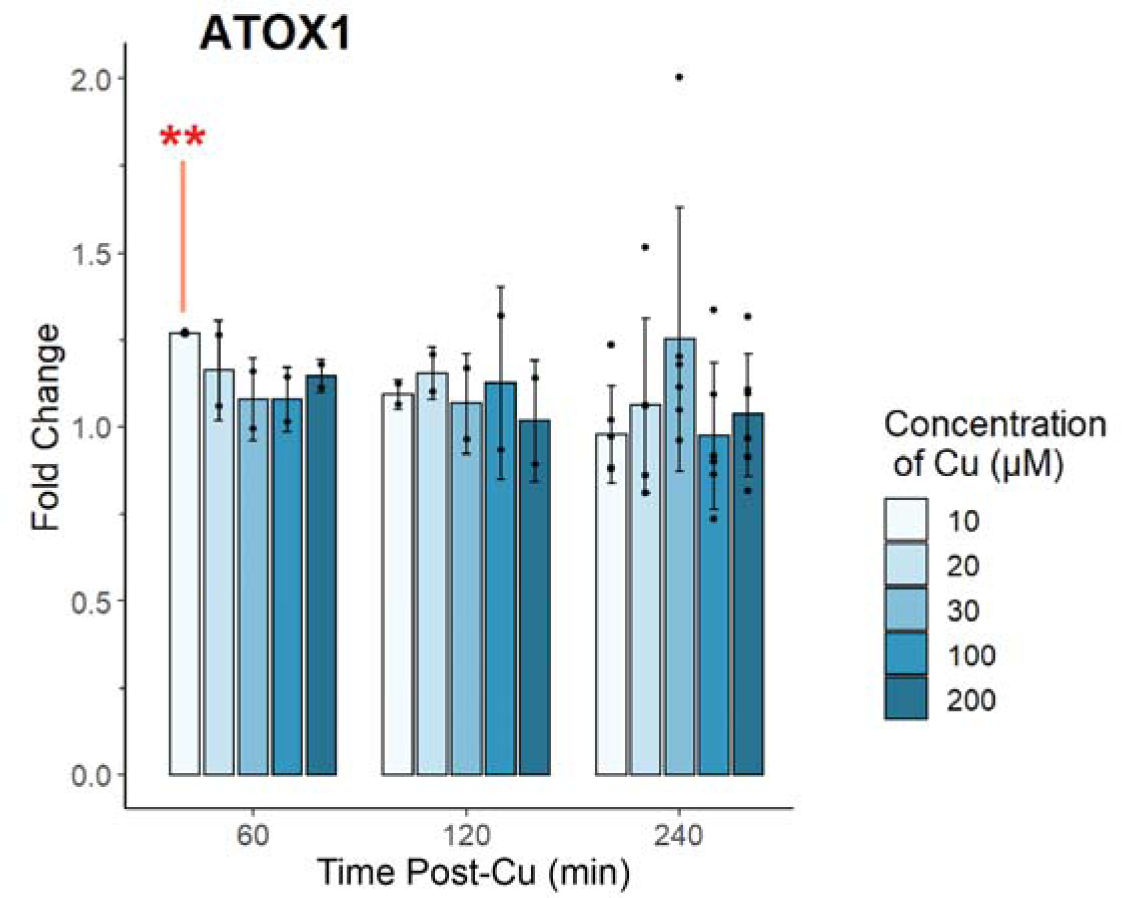
Copper-stimulated changes in A549 *ATOX1* expression levels. RNA-qPCR results of RNA extracted from A549 cells following supplementation with a screen of CuCl_2_ concentrations for the times indicated on the X-axis. As expected, there were no significant changes following CuCl_2_-supplementation, except for the one anomalous condition after 60 minutes at the lowest CuCl_2_ supplementation level. Individual datapoints correspond to biological replicates. Error bars denote standard deviation. *n* = 2, 2, 6, for 60 min, 120 min, and 240 min post-Cu, respectively.

**Fig. S5.**
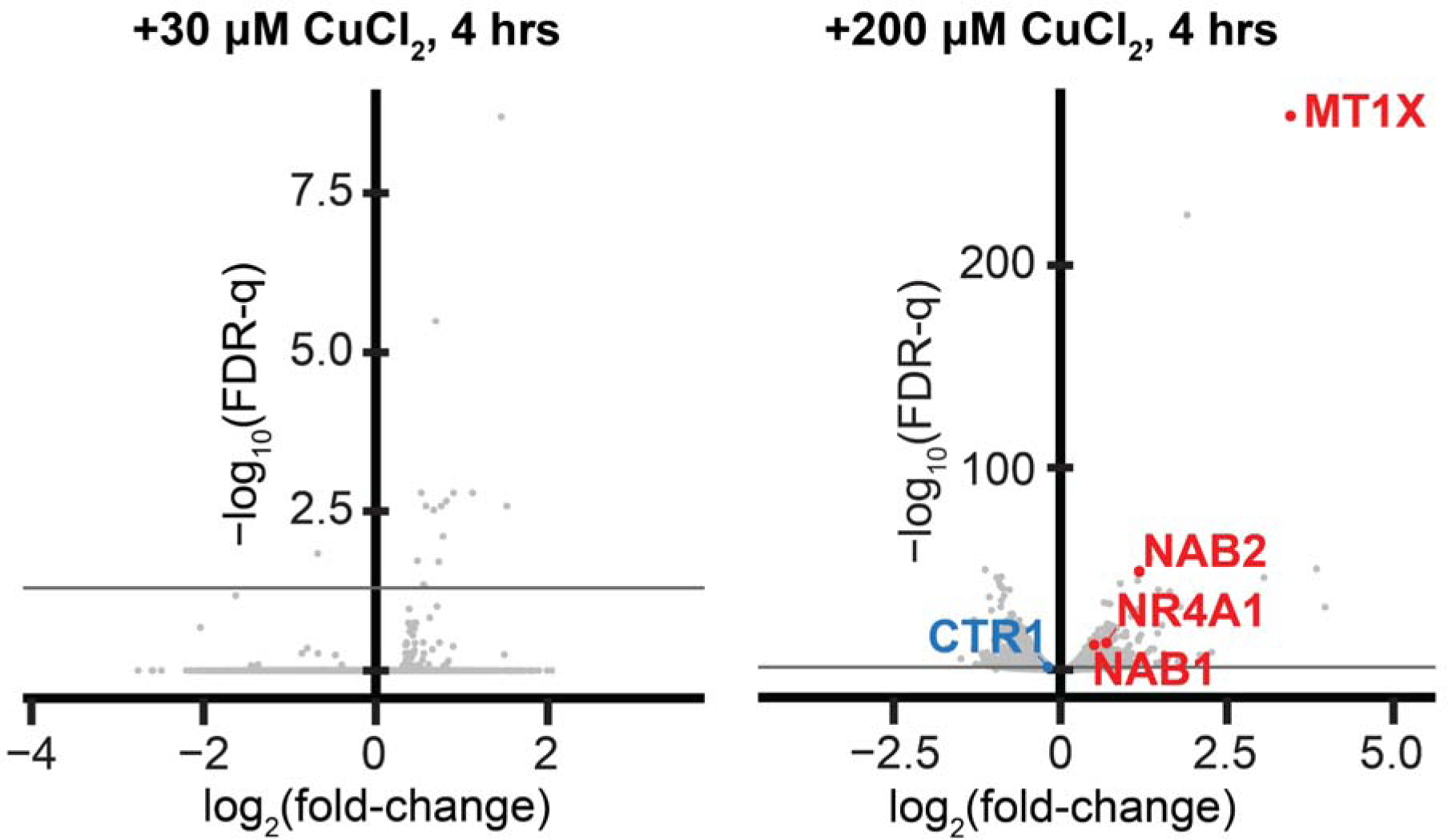
Copper-supplemented A549 RNA-seq volcano plots. Plots were generated from cells supplemented with CuCl_2_ for 4 hours relative to unsupplemented cells. Select transcripts with significantly altered expression patterns post-Cu treatment are labeled with standard nomenclature (blue for downregulated, red for increased expression). No transcripts of interest were significantly altered by 30 μM CuCl_2_ treatment for 4 hours. Gray horizontal line denotes FDR-q = 0.05. The decrease in *CTR1* expression levels 4 hours post-supplementation with 200 μM CuCl_2_ is statistically significant (FDR-q < 0.05). *n* = 3 biological replicates for each condition.

**Fig. S6.**
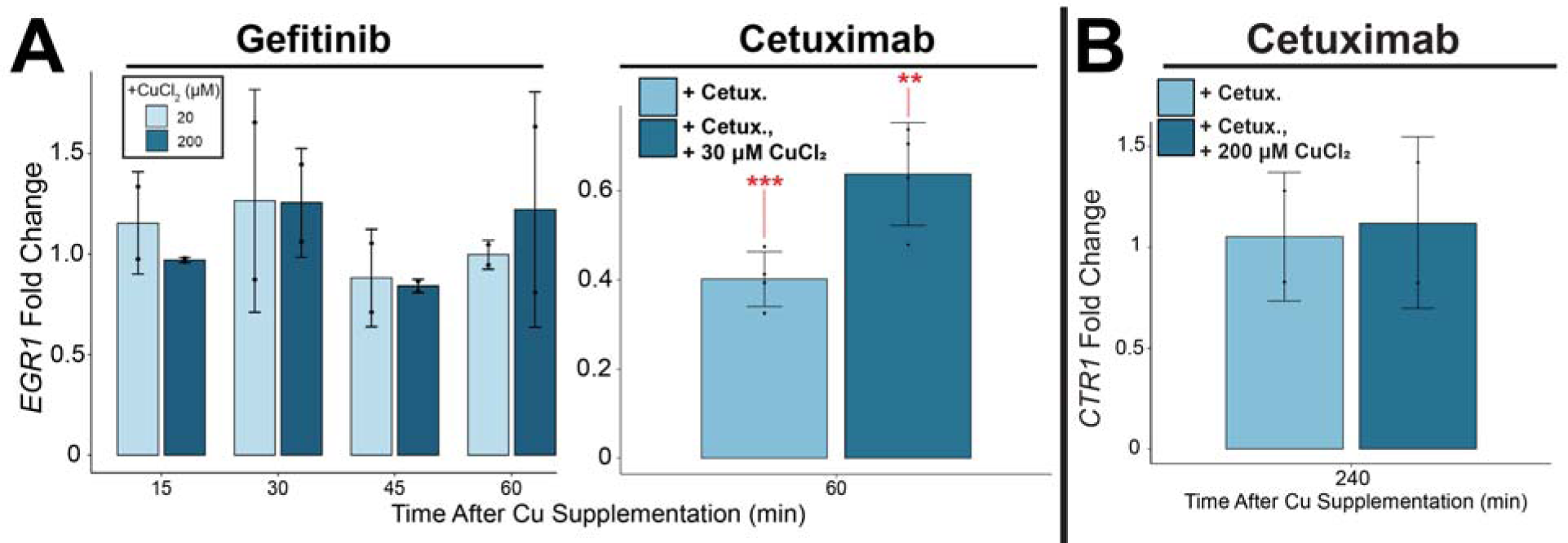
Effects of EGFR inhibitors on copper-stimulated changes in *EGR1* and *CTR1* expression levels. RNA-qPCR of (*A*) *EGR1* and (*B*) *CTR1* transcript levels following EGFR inhibitor pre-treatment, and subsequent CuCl_2_ or no addition (control). Data points correspond to biological replicates. *EGR1* levels were unaltered following copper-treatment of gefitinib (an EGFR inhibitor compound)-pretreated cells and were lower in copper-treated, cetuximab (an EGFR inhibitor monoclonal antibody)-pretreated cells relative to untreated cells. *CTR1* levels, conversely, were unchanged in cetuximab pre-treated cells supplemented with CuCl_2_. Collectively, these results validate copper-driven activation of MAPK/ERK signaling and associated transcriptional activities, as well as copper-stimulated *CTR1* transcriptional repression. Error bars denote standard deviation. *n* = 2, 4, and 2 biological replicates for gefitinib pretreated *EGR1* fold-change, Cetuximab pre-treated *EGR1* fold-change, and Cetuximab pre-treated *CTR1* fold-change.

**Fig. S7.**
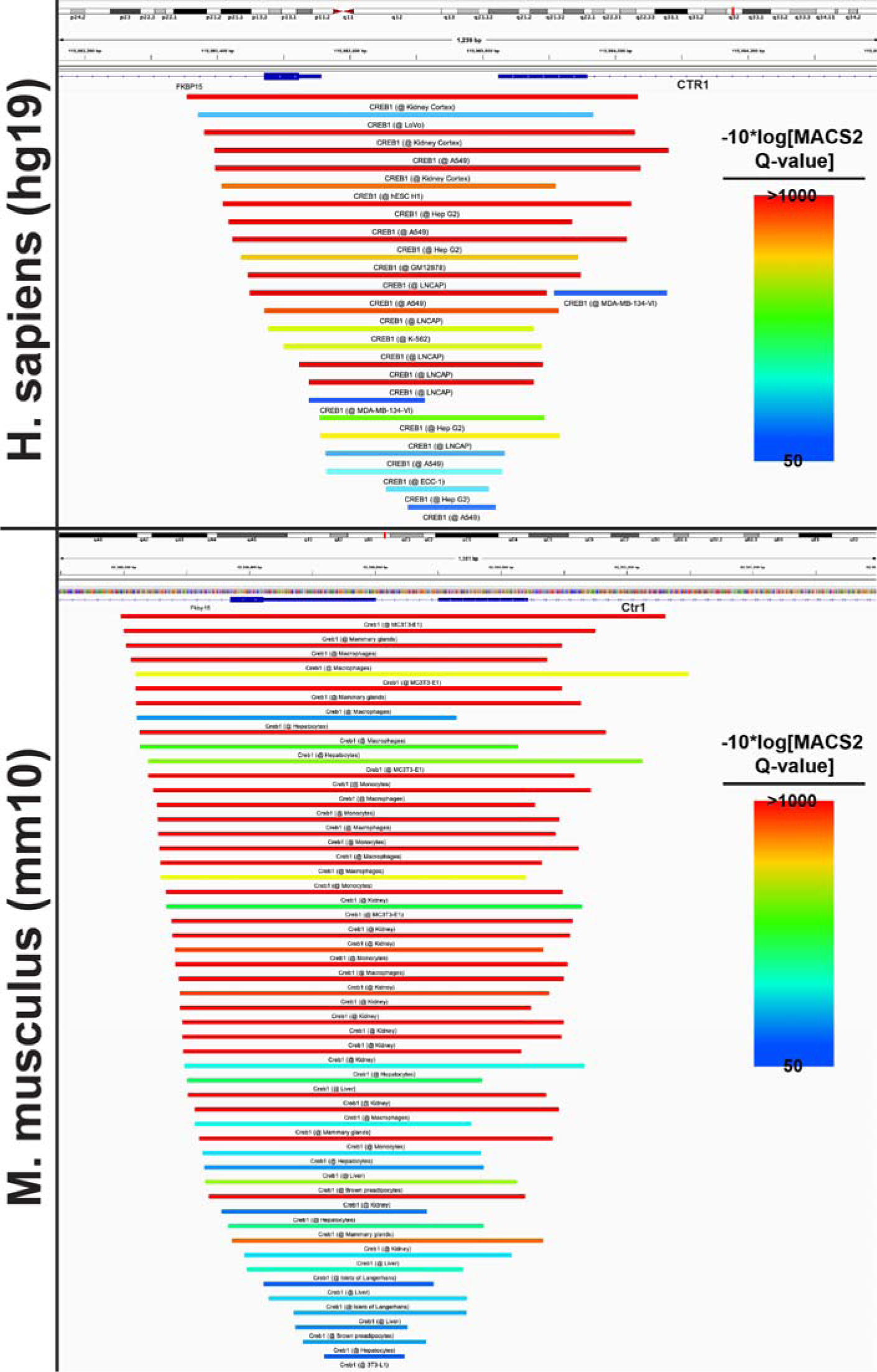
All CREB1 ChIP-Atlas results. Bed file summary of all *Homo sapiens* (genome build hg19) and *Mus musculus* (genome build mm10) CREB1/Creb1 ChIP-seq experiments in the ChIP-Atlas, corresponding to CREB1 binding in the *CTR1*/*Ctr1* promoter. 22 of 31 total human and 52 of 81 total mouse ChIP-seq experiments identified significant binding in the *CTR1*/*Ctr1* promoter, as shown above. MACS2 Q-value denotes the FDR-adjusted *p*-value.

**Fig. S8.**
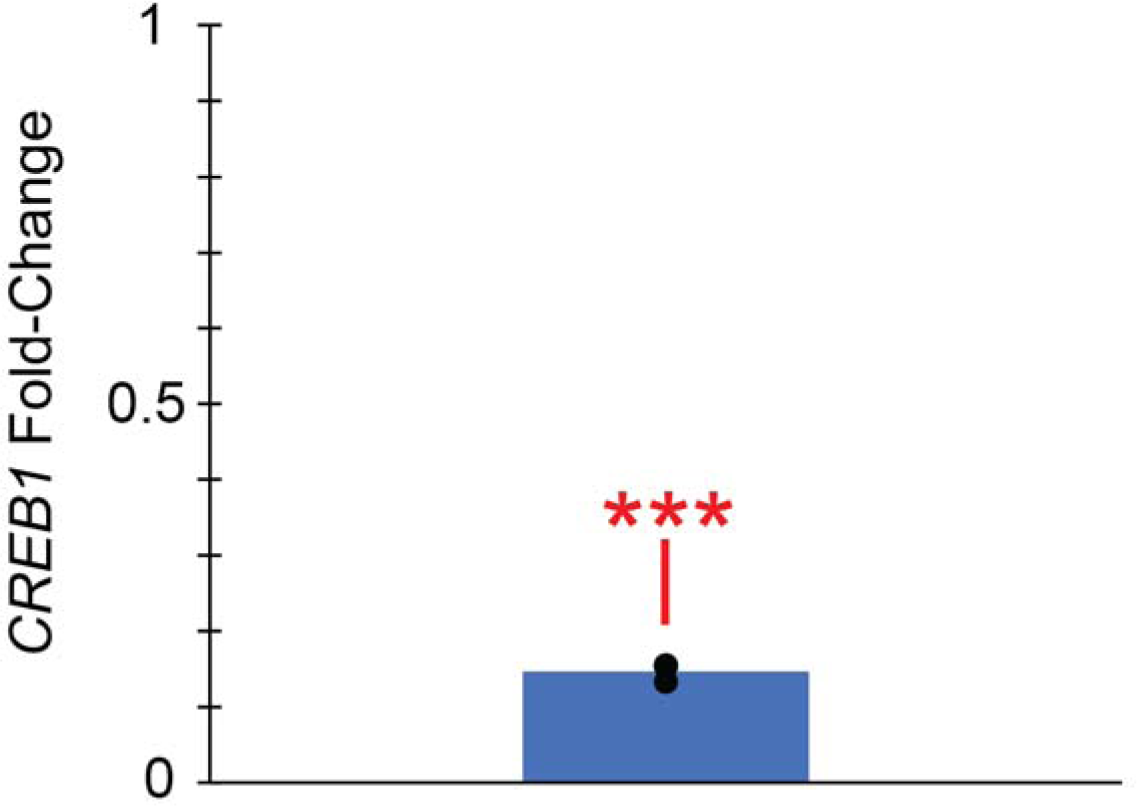
*CREB1* transcript expression fold-change following CREB1 siRNA knockdown in A549 cells (relative to negative control siRNA transfection) for *n* = 3 biological replicates.

**Fig. S9.**
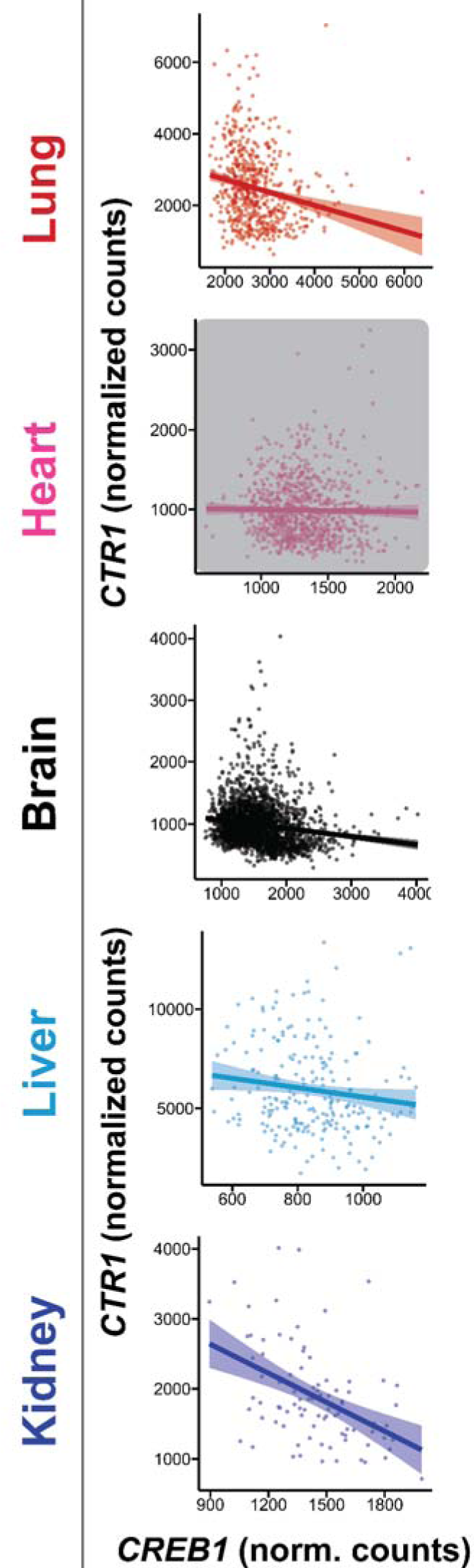
*CTR1*/*CREB1* RNA-seq expression value correlations. Correlation of *CTR1* transcript expression levels with those of *CREB1* from GTEx RNA-seq datasets across the five vital organs. The statistically insignificant correlation in the heart tissue is covered by a gray box. Sample sizes are the same is in Fig. 3.

**Fig. S10.**
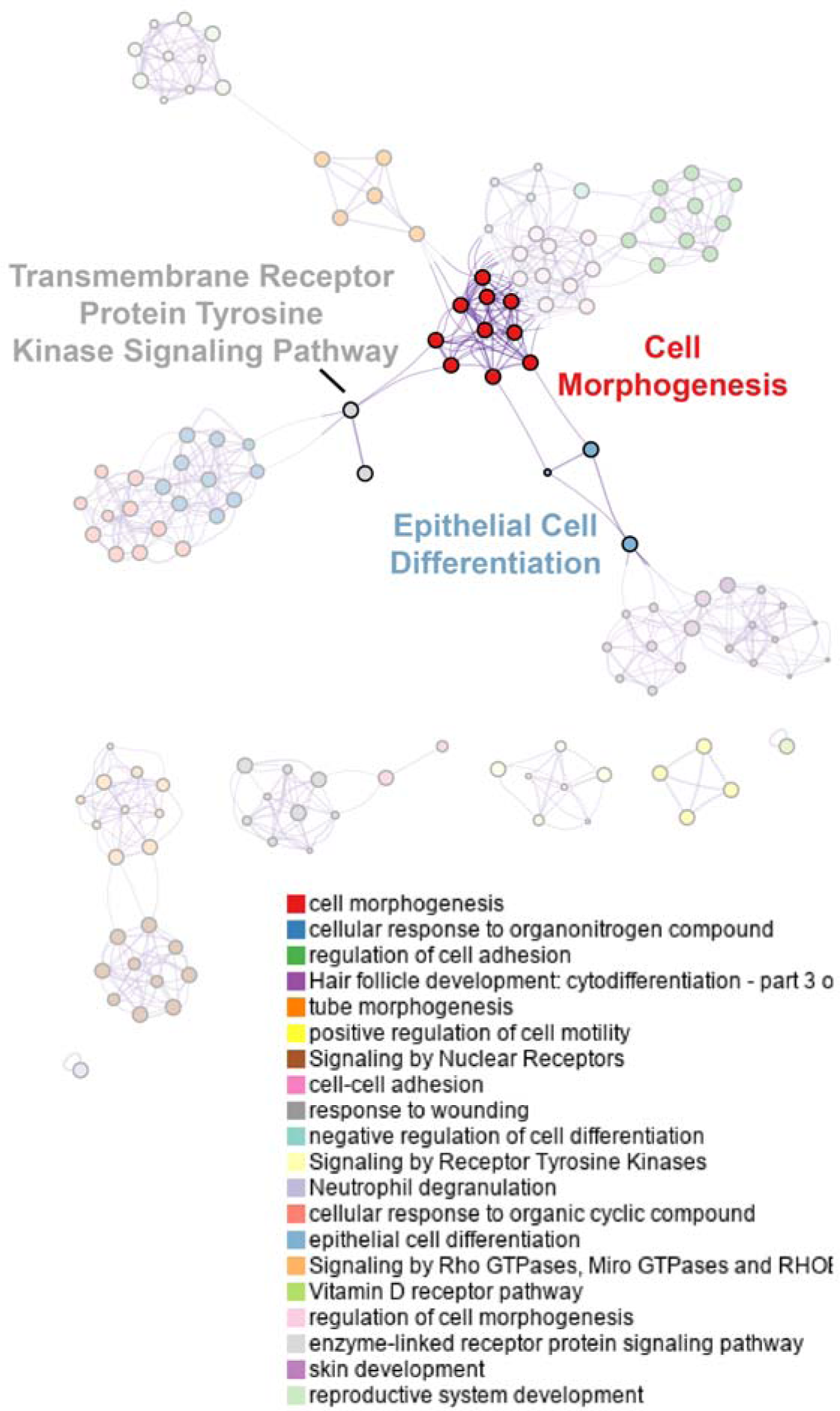
Enriched pathways/process network generated from transcripts significantly downregulated by TM treatment of MDA-MB-468 cells.

**Fig. S11.**
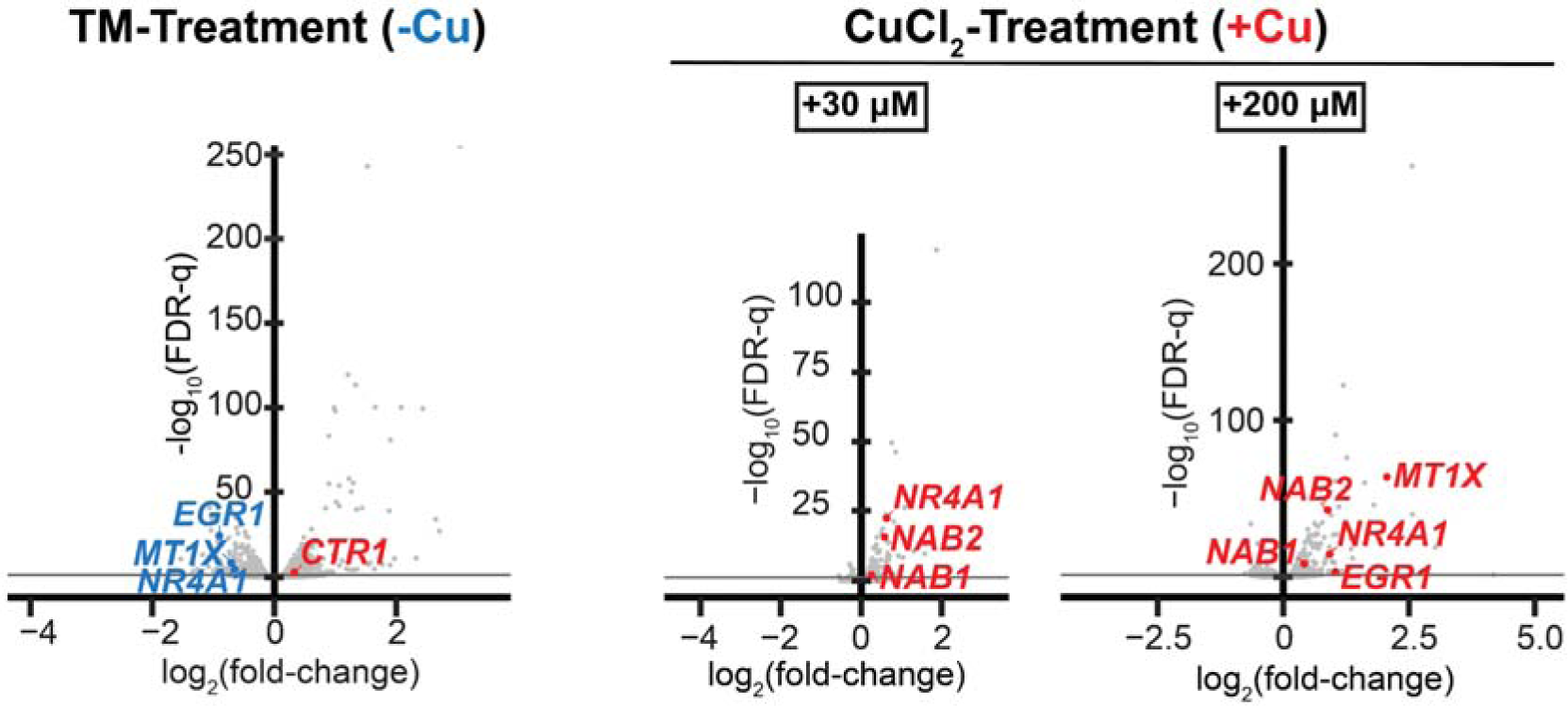
Cu- or TM-supplemented MDA-MB-468 RNA-seq volcano plots. Full volcano plots depicting differential expression results from TM-treated MDA-MB-468 RNA-seq, as well as CuCl_2_-treated A549 cells for 2 hours, as featured in Fig. 2. Gray horizontal line denotes FDR-q =0.05. *n* = 2 for TM, 3 for Cu: biological replicates for each condition.

**Fig. S12.**
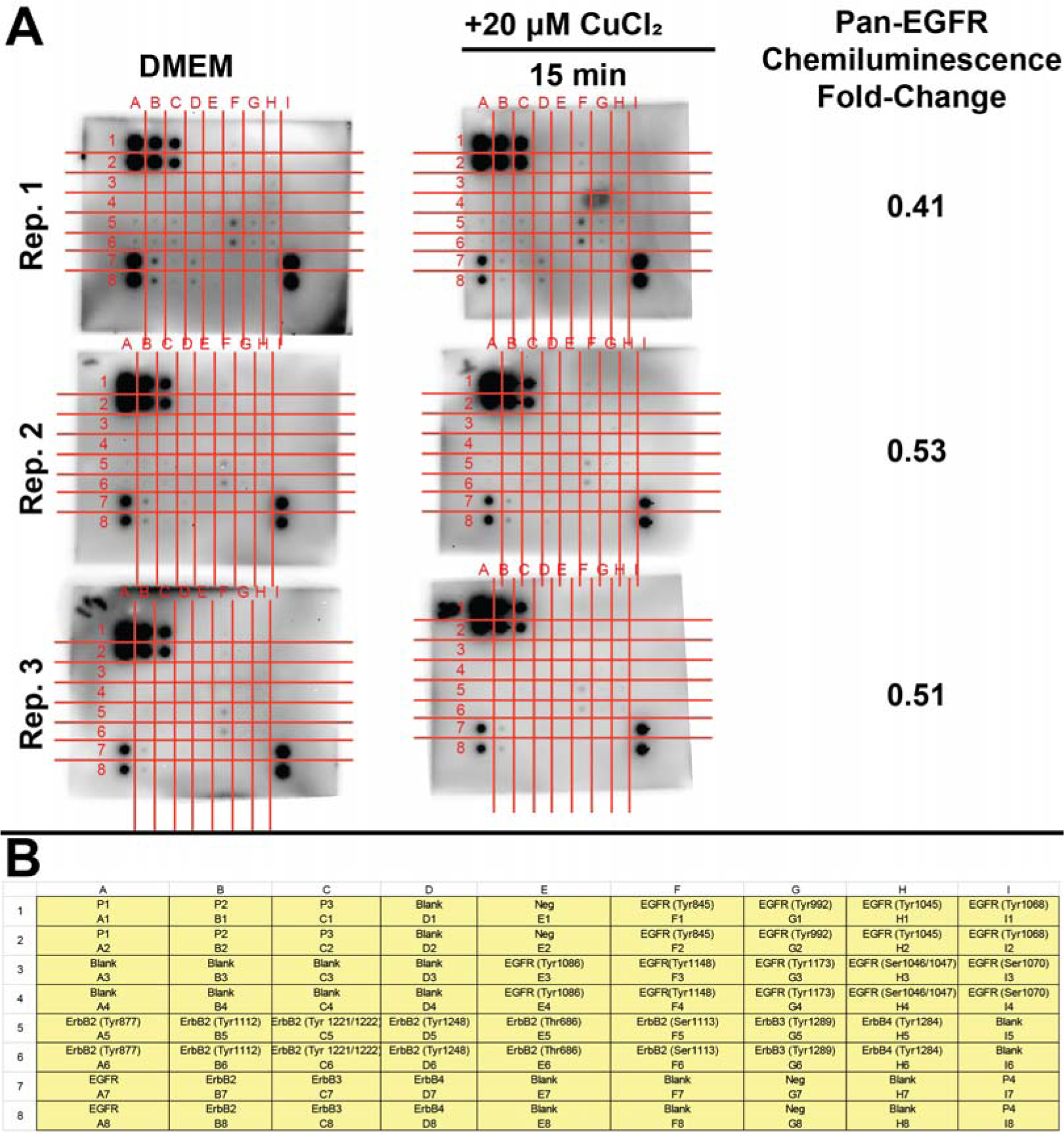
EGFR phosphorylation antibody arrays. (*A*) EGFR-phosphorylation antibody arrays, with change in pan-EGFR chemiluminescence intensity (as measured in ImageJ) comparing CuCl_2_ supplemented to unsupplemented lysates of the same biological replicate. In practice, subtle differences in chemiluminescence intensity for the EGFR phosphorylation sites were difficult to detect against the highly variable intensity of the membrane array background. (*B*) Key denoting antibody location identities from the arrays. P1-4 are reference (control) antibody spots. *n* = 3 biological replicates for each condition.

**Fig. S13.**
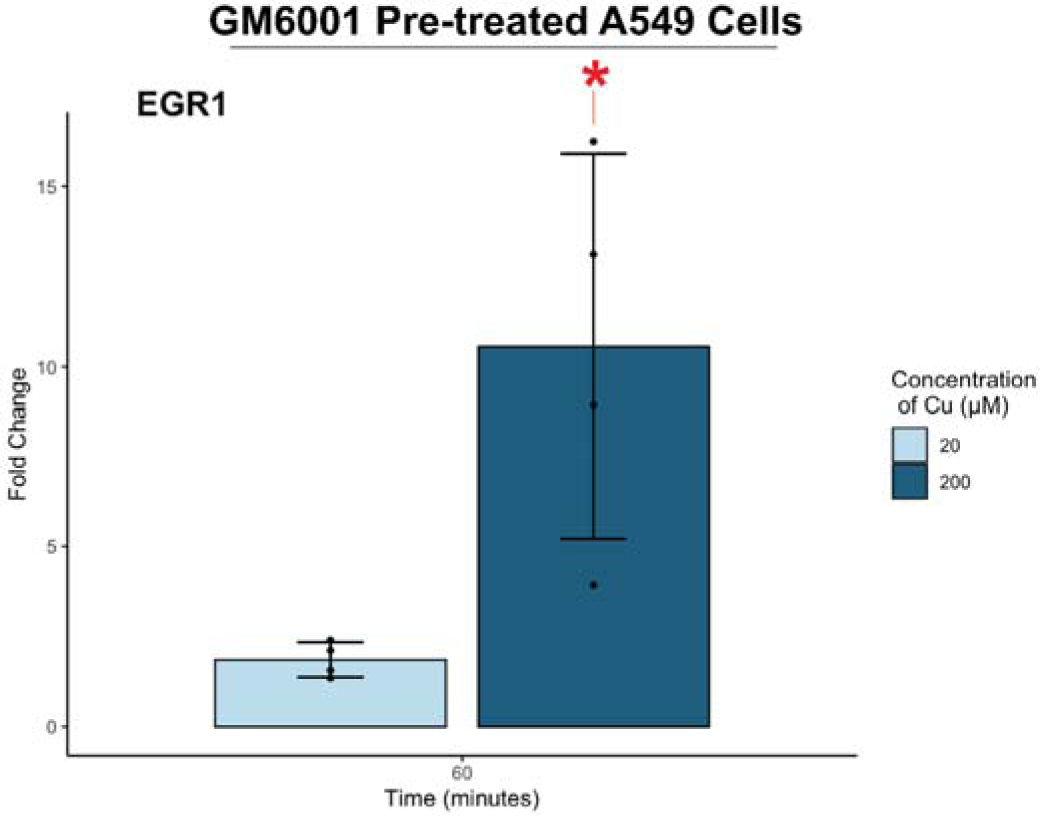
Effect of metalloproteinase inhibitor pretreatment on copper-stimulated changes in *EGR1* expression levels. RNA-qPCR detected changes in *EGR1* transcript levels in RNA extracted from A549 cells following pretreatment with 20 µM GM6001 for 30 minutes and then CuCl_2_ treatment for 60 minutes (relative to cells not supplemented with CuCl_2_). Error bars denote standard deviation. *n* = 4 biological replicates for each condition.

**Fig. S14.**
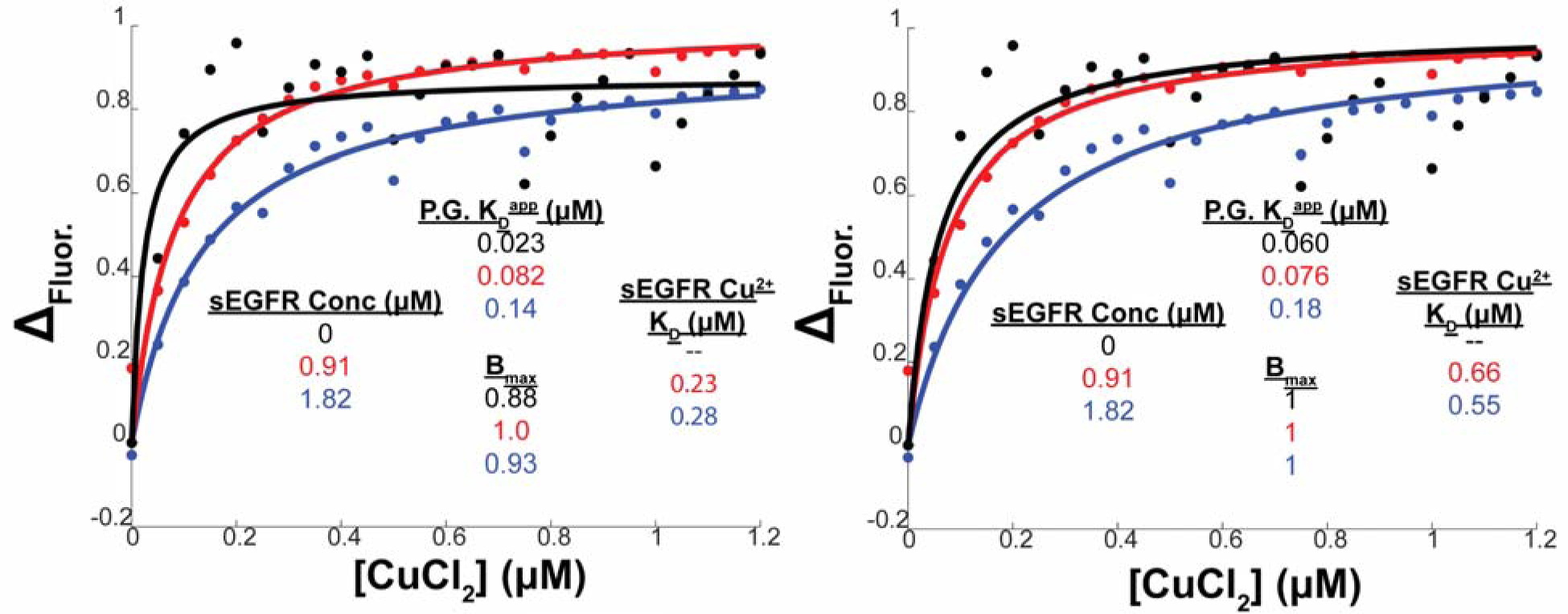
sEGFR and Phen Green Cu^2+^-binding competition. Fit to Phen Green Cu^2+^-binding competition, (***left***) allowing B_max_ to float through the fitting procedure or **(*right***) fixing B_max_ at 1. See materials and methods for definitions of variables and methods for calculations. Importantly, we measured a Phen Green-Cu^2+^ K_D_ of 0.023-0.060 μM, very close to the previously reported value of 0.015 μM^67^. Definitions of affinity terms provided in the Materials and Methods.

**Fig. S15.**
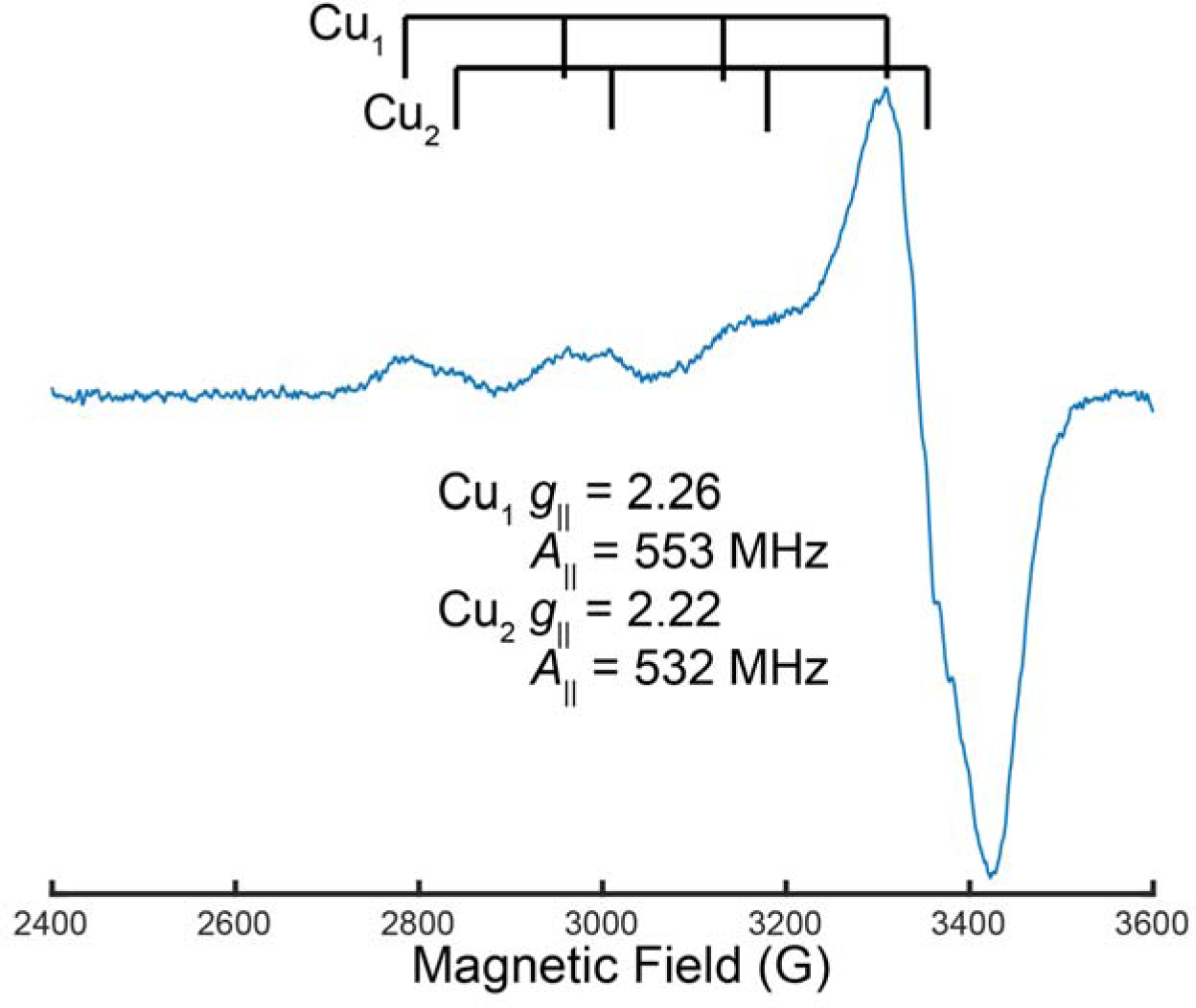
X-band EPR spectrum of sEGFR loaded with 1 Cu^2+^ equivalent. The *g*_||_ regions of the spectrum defines two well-structured Cu^2+^ coordination spheres, corresponding to either two distinct sites or one site with two slightly different ligation spheres. Collection conditions were as follows; 9.631 GHz microwave frequency, 30 dB microwave power attenuation, 9 G modulation amplitude, 100 kHz modulation frequency, 10.24 ms time constant, 84 s scan rate, 20 K temperature. Brackets denote the Cu^2+^ hyperfine splitting *A*_||_.

**Fig. S16.**
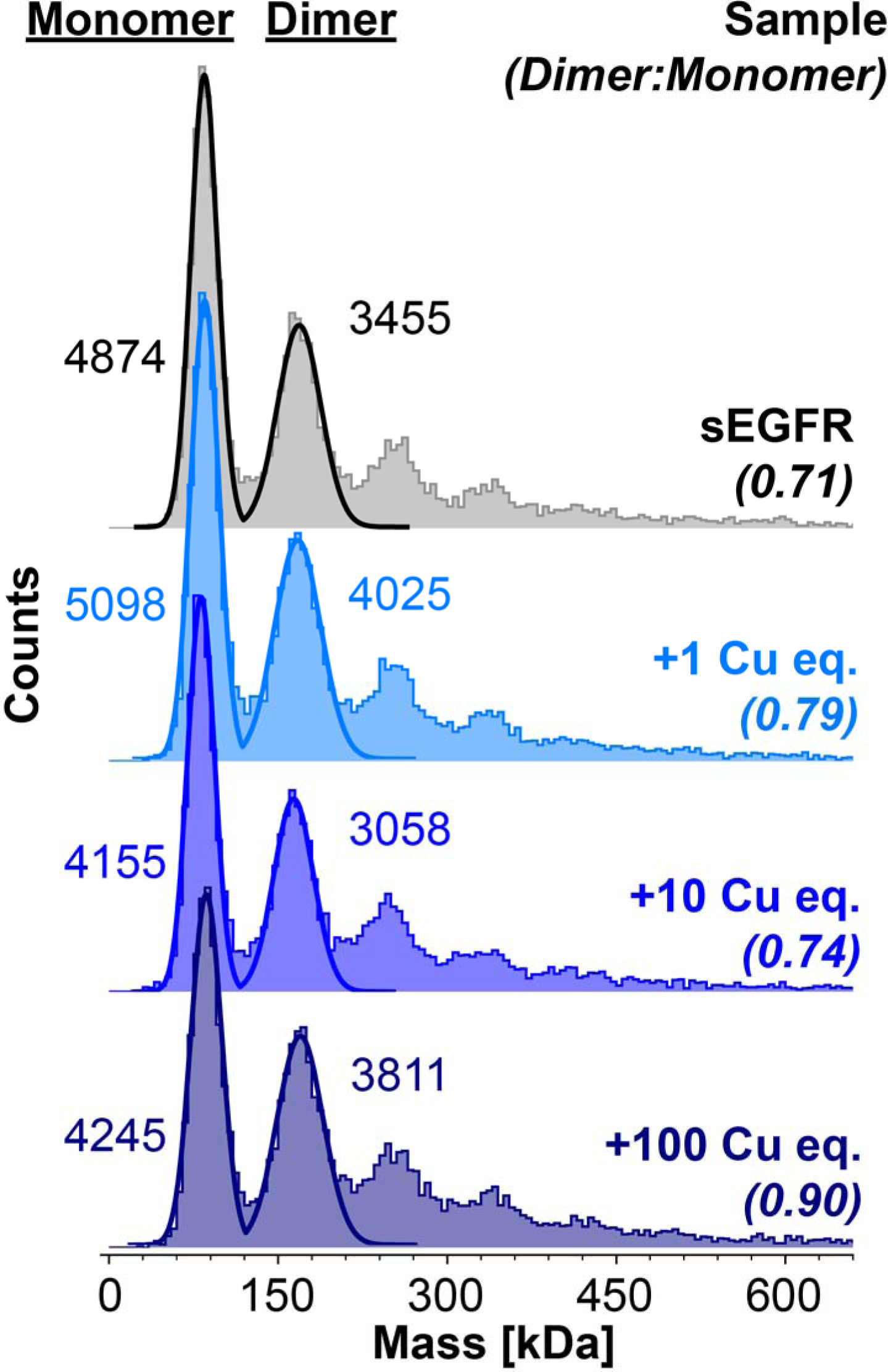
Mass photometry of sEGFR (100 nM) incubated overnight (4°C) with various CuCl_2_ equivalents. As the starting concentration of sEGFR was 100 nM, the 100 equivalent loaded sample corresponds to 10 μM Cu^2+^, well above the experimentally-determined K_D_ (200-700 nM) of sEGFR for Cu^2+^.

**Fig. S17.** Proteome Profiler Antibody Arrays for HEK 293T cells treated with ZnCl_2_ (20 μM). The layout (and antibody spot locations) is the same as depicted in SI Appendix Fig. S3.

**Fig. S18.**
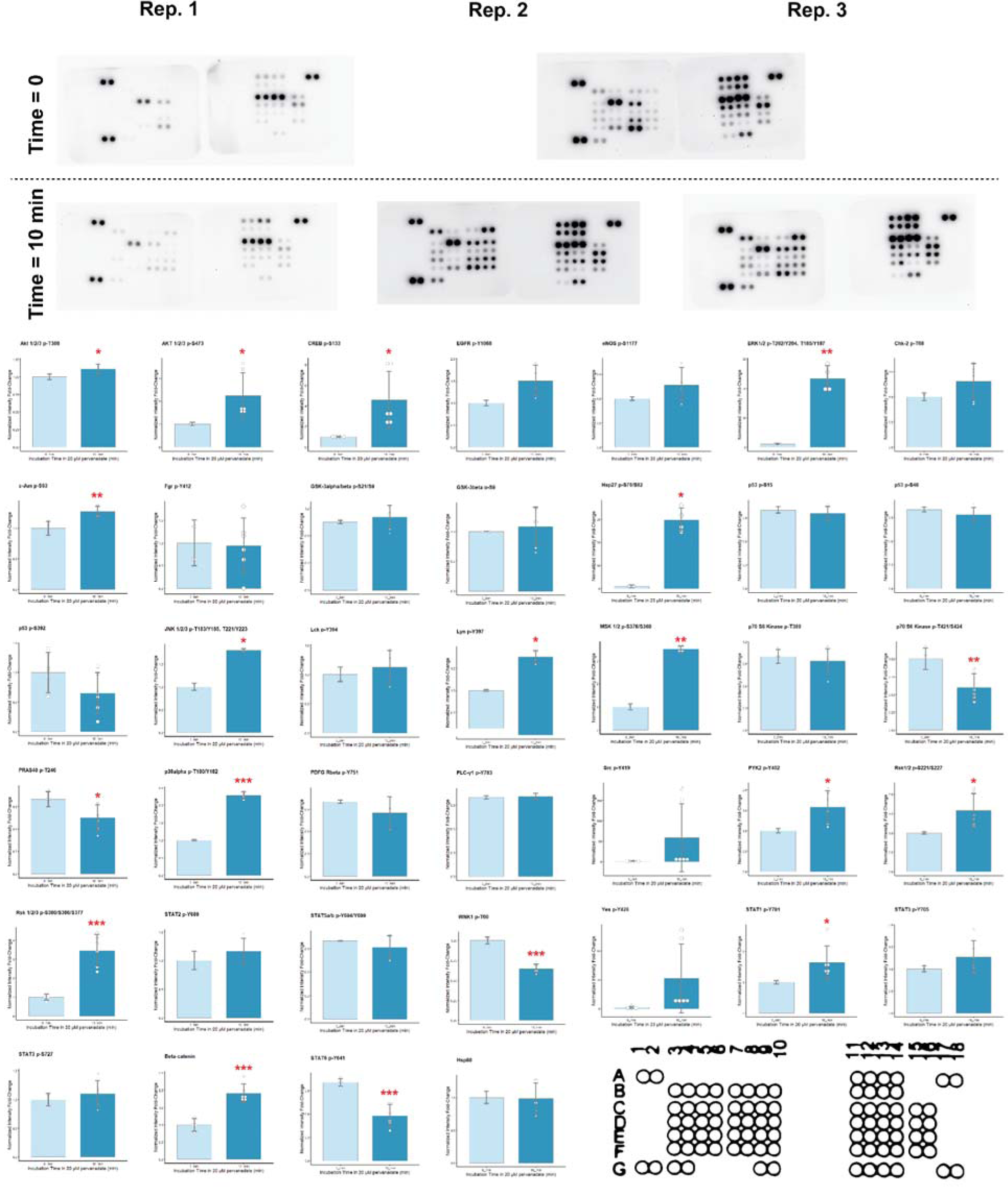
Proteome Profiler Antibody Arrays for HEK 293T cells treated with pervanadate (20 μM). The layout (and antibody spot locations) is the same as depicted in SI Appendix Fig. S3.

**Fig. S19.**
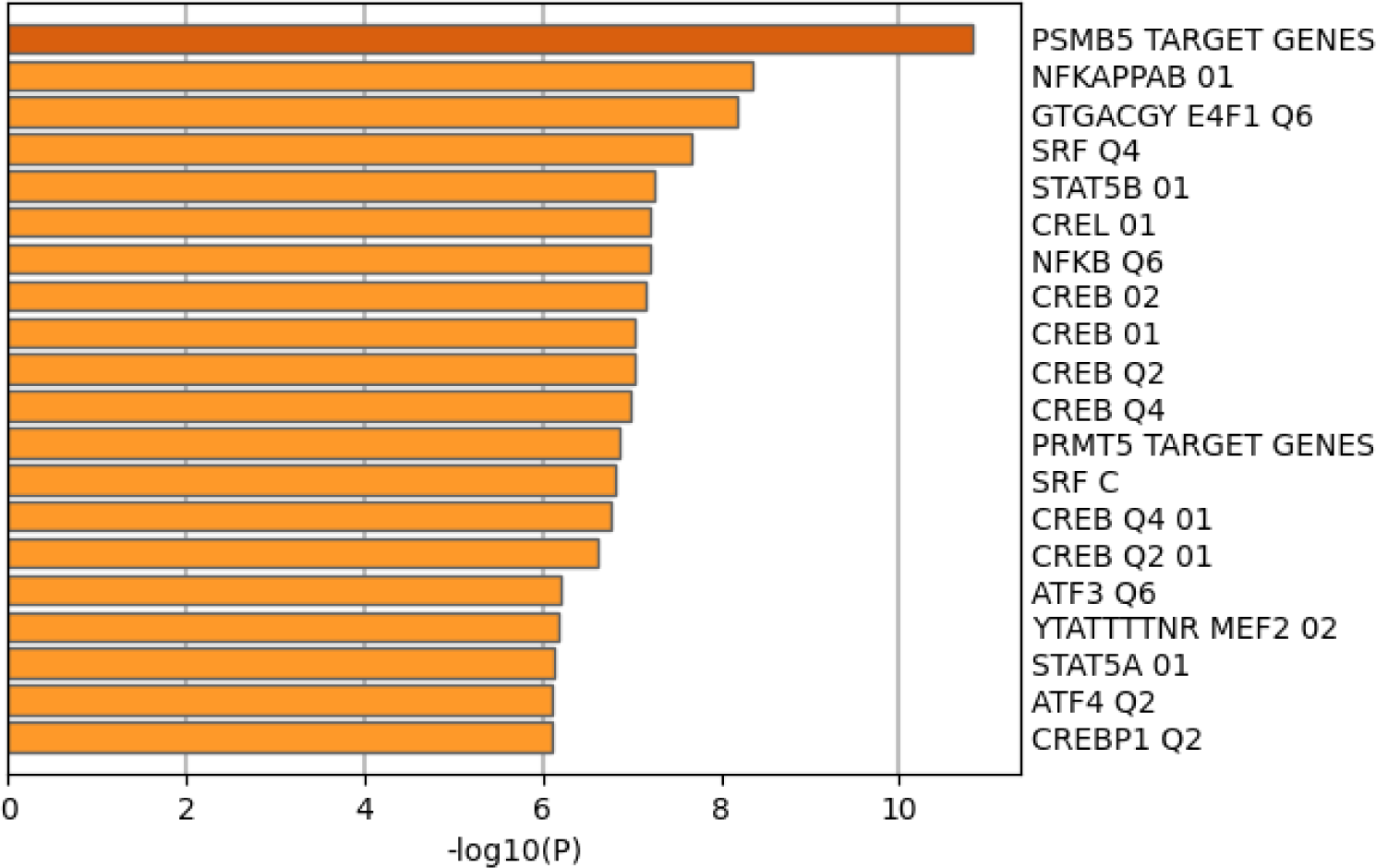
Fully-labelled TF enrichment generated from significantly differentially expressed transcript post-30 µM CuCl_2_ treatment.

**Fig. S20.**
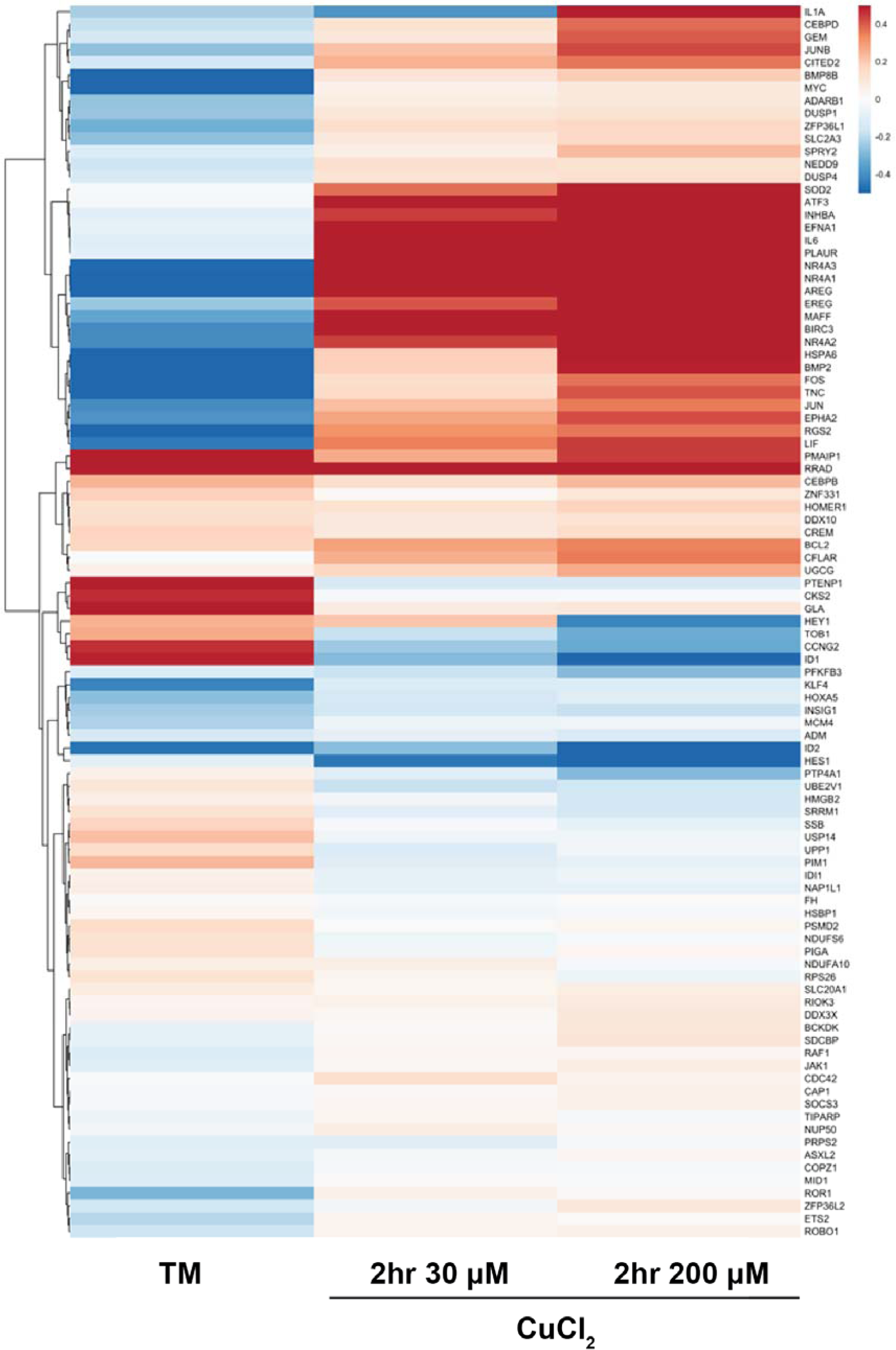
CREB1-target gene hierarchical cluster. Hierarchical cluster of differential expression of CREB1-target genes in the TM-treated and 2 hr CuCl_2_-treated RNA-seq results. List of CREB1-target genes was generated from the human cell line cAMP-responsive genes in ^34^ – of the 195 human cAMP-responsive genes, 97 produced responses following DESeq2 analysis in TM and copper conditions, as shown in the hierarchical cluster above.

**Table S1.** CREB Target Gene Database results for human *SLC31A1* (*CTR1*) (accessed 6/18/22). CREB or p-CREB (phospho-CREB) indicates the ChIP target for ChIP-seq experimental results reported below. “0h”, “1h”, and “4h” for HEK samples indicate duration of forskolin stimulation (to generate cAMP). CRE prediction value “CRE_NoTATA” indicates a predicted functional CRE without the presence of a proximal TATA box.

| Target Gene | HEK CREB<br>(Binding Ratio, *p*-value) | Hepatocyte CREB<br>(Binding Ratio, *p*-value) | Hepatocyte p-CREB<br>(Binding Ratio, *p*-value) | Islet p-CREB<br>(Binding Ratio, *p*-value) | HEK p-CREB<br>0h<br>1h<br>4h<br>(Binding Ratio, *p*-value) | CRE prediction |
| --- | --- | --- | --- | --- | --- | --- |
| *CTR1* | 3.8, 2.50E-09 | 2.1, 4.80E-05 | 3.6, 5.60E-07 | 1.1, 1.50E-01 | 2.6, 3.50E-07<br>3.9, 3.60E-11<br>3.5, 6.40E-09 | CRE_NoTATA |

**Table S2.** KnockTF results assessing *SLC31A1*/*CTR1* expression level changes induced by CREB1 knockdown. Statistically-significant (FDR-q < 0.05) change measured in K562 cells shown in yellow, with statistically-insignificant results shown in gray.

| Target Gene | TF | Knock-Method | Tissue Type | Biosample | Fold-Change | p-value | FDR-q | Dataset ID |
| --- | --- | --- | --- | --- | --- | --- | --- | --- |
| *CTR1* | CREB1 | shRNA | Haematopoietic and lymphoid tissue | K562 | 1.55975 | 1.46E-07 | 2.45E-05 | DataSet_01_004 |
| *CTR1* | CREB1 | shRNA | Parotid gland | H3118 MEC | 0.60725 | 1.01E-02 | 1.84E-01 | DataSet_01_194 |
| *CTR1* | CREB1 | shRNA | Parotid gland | HSY | 0.61149 | 5.12E-03 | 3.20E-01 | DataSet_01_195 |

**Table S3.**
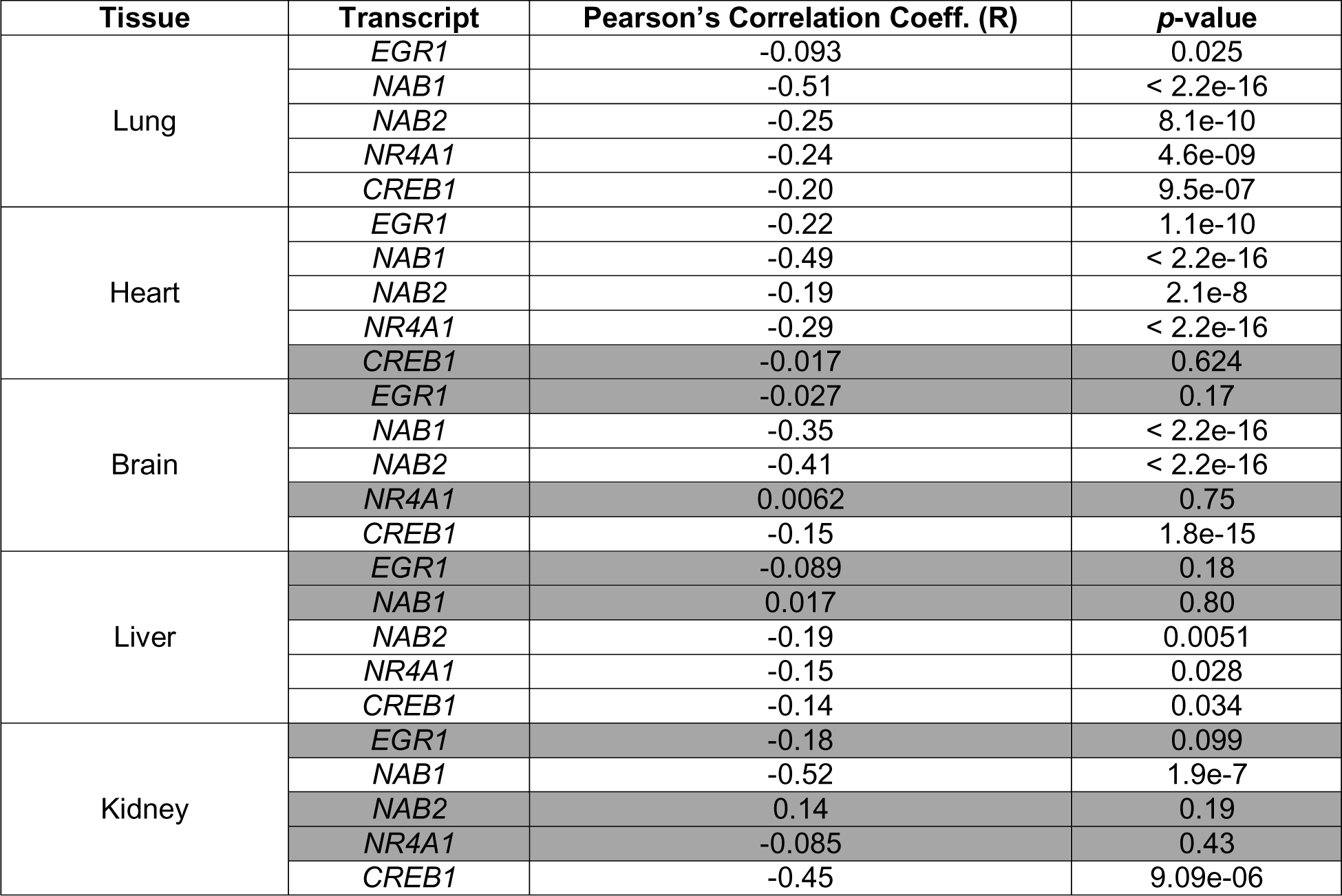
GTEx transcript expression level correlations with *CTR1* expression levels. Statistically-insignificant correlations shown in gray.

**Table S6.**
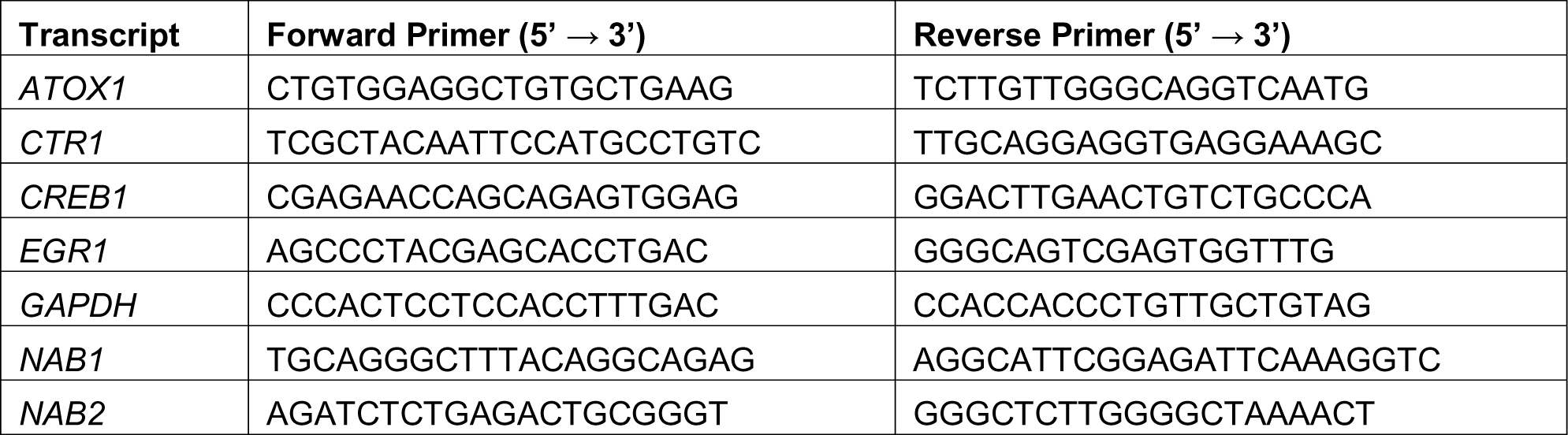
Primers used for RNA-qPCR experiments.

